# Genomic stability in the Galápagos *Scalesia* adaptive radiation: Consistent transposable element accumulation despite hybridization and ecological niche shifts

**DOI:** 10.1101/2024.09.30.614436

**Authors:** José Cerca, Patricia Jaramillo Díaz, Clément Goubert, Heidi Yang, Vanessa C. Bieker, Mario Fernández-Mazuecos, Pablo Vargas, Rowan Schley, Siyu Li, Juan Ernesto Guevara-Andino, Bent Petersen, Gitte Petersen, Neelima R. Sinha, Lene R. Nielsen, James H. Leebens-Mack, Gonzalo Rivas-Torres, Loren H. Rieseberg, Michael D. Martin

**Affiliations:** Department of Natural History, University Museum, Norwegian University of Science and Technology (NTNU), Trondheim, Norway; Centre for Ecological and Evolutionary Synthesis (CEES), Department of Biosciences, University of Oslo, Oslo, Norway; Estación Científica Charles Darwin, Fundación Charles Darwin, Santa Cruz, Galápagos, Ecuador; Department of Botany and Plant Physiology, University of Málaga, Málaga, Spain; IUCN SSC Galapagos Plant Specialist Group, Puerto Ayora, Galapagos 200102, Ecuador; McGill Genome Centre, McGill University, Montreal H3A 0G1, QC, Canada; Department of Ecology & Evolutionary Biology, University of California, Los Angeles, Los Angeles, California, United States of America; Departamento de Biología (Botánica), Facultad de Ciencias, Universidad Autónoma de Madrid, Calle Darwin 2, 28049 Madrid, Spain; Centro de Investigación en Biodiversidad y Cambio Global, Universidad Autónoma de Madrid (CIBC-UAM), Calle Darwin 2, 28049 Madrid, Spain; Biodiversity and Conservation, Real Jardín Botánico, 28014 Madrid, Spain; Department of Geography, University of Exeter, Laver Building, North Park Road, Exeter, Devon, UK; Department of Plant Biology, University of California, Davis, Davis, CA, 95616, USA; Grupo de Investigación en Ecología y Evolución en los Trópicos-EETrop- Universidad de las Américas, Quito-Ecuador; Center for Evolutionary Hologenomics, Globe Institute, University of Copenhagen, DK-1353, Copenhagen, Denmark; Centre of Excellence for Omics-Driven Computational Biodiscovery (COMBio), Faculty of Applied Sciences, AIMST University, Kedah, Malaysia; Department of Ecology, Environment and Plant Sciences, Stockholm University, 106 91 Stockholm, Sweden; Department of Geosciences and Natural Resource Management, University of Copenhagen, Rolighedsvej 23, 1958 Frederiksberg C, Denmark; Department of Plant Biology, University of Georgia, Athens, GA, 30602, USA; Colegio de Ciencias Biológicas y Ambientales, Galapagos Science Center, Universidad San Francisco de Quito USFQ, Quito 170901, Ecuador; Department of Botany and Biodiversity Research Centre, University of British Columbia, Vancouver, BC, V6T 1Z4, Canada

## Abstract

Transposable elements (TEs) have been hypothesized to play a pivotal role in driving diversification by facilitating the emergence of novel phenotypes and the accumulation of divergence between species. Hybridization and adaptation to novel niches are hypothesized to influence TE accumulation in genomes by disrupting genomic regulation, including the suppression of TE replication. The rapid speciation and ecological diversification characteristic of adaptive radiations offer a unique opportunity to examine the link between TE accumulation and speciation, diversification, hybridization and adaptation. Here, focusing on all 15 species of the genus *Scalesia* (Asteraceae), a radiation endemic to the Galápagos Islands, we test whether hybridization or shifts in ecological niche are associated with changes in TE accumulation in genomes. Our analyses reveal little to no variation in TE accumulation among *Scalesia* species nor its hybrid populations. Shifts in ecological niches, linked to climatic variation, did not result in discernible changes in TE accumulation, a surprising finding given the anticipated selective pressure imposed by aridity, a factor often linked to genome size reduction. We found no distinct patterns in the temporal accumulation of TEs, and no effects at the class or superfamily level. Our findings challenge the assertion that TEs have directly driven diversification, speciation and local adaptation. Instead, we suggest that TEs may simply be ‘along for the ride,’ rather than actively contributing to plant diversification.

## Introduction

Transposable elements are DNA sequences capable of relocating within the genome [1]. Through their mobility and ability to replicate, TEs are able to generate a diverse range of mutations encompassing changes in promoter gene sequences, frameshifts, intronic modifications and gene duplications within their hosts [2–8]. Some of these mutations have been linked to the evolution of novel phenotypes. For instance, the evolution of the melanic phenotype of the peppered moth (*Biston betularia*) showcases the potential functional consequences of TE insertions as the insertion of the ‘carb-TE’ in an intronic region of the *cortex* gene alters its expression, likely contributing to the melanic phenotype [9].

Advances in genomic sequencing technology and processing algorithms have greatly facilitated the detection and characterization of TEs, particularly in non-model organisms with limited genomic resources [10]. The evident adaptive potential of TEs in shaping phenotypes [4,11,12], along with the evidence of substantial variation in TE diversity and accumulation across different reference genomes (e.g. [13,14], has intensified interest in a potential relationship between TEs and diversification. One particularly intriguing hypothesis is that lineages with high diversification rates also exhibit elevated rates of TE accumulation on their genomes [6,15–22]. A key limitation of these studies has been their reliance on a small number of genomes, often with limited taxonomic coverage while encompassing large clades, and primarily based on correlations. Consequently, there may be an overestimation of the role of TEs as major drivers of diversification at macroevolutionary scales.

Adaptive radiations represent evolutionary experiments where extensive ecological variation aligns with rapid speciation, and they have been a cornerstone of our understanding of evolutionary change [23– 25]. Adaptive radiations can also yield valuable insights into the interplay between TEs and evolutionary success; however, the evidence available is conflicting and largely originated from cichlid radiations. For example, a study examining four species within an African cichlid adaptive radiation found higher TE counts in the genomes and transcriptomes of the radiating species compared to non-radiating lineages [26]. More recent evidence, however, suggested a lack of correlation between species richness at the tribe level in the Lake Tanganyika adaptive radiation and TE content across 245 genomes [27]. Similarly, cichlid lineages in the Americas had no discernible differences in TE content when contrasting a radiating lineage (*Amphilophus citrinellus*) with a non-radiating lineage (*Archocentrus centrarchus*; [28]. Notably, a golden phenotype in the cichlid genus *Amphilophus* is associated with a transposon-derived inverted repeat in the *goldentouch* gene [11]. This conflicting body of evidence underscores the necessity for further exploration of TEs as facilitators of diversifications, particularly extending beyond the scope of cichlid radiations.

The *Scalesia* radiation (Asteraceae, Asterales, Magnoliidae), endemic to the Galápagos archipelago, presents an excellent opportunity to investigate the potential relationship between TEs and diversification. This genus comprises ∼15 species that evolved from a single common ancestor which diversified about 1 million years ago [29]. The recent origin of this radiation may facilitate the reconstruction and tracking of TE accumulation over time. Furthermore, *Scalesia* exhibits remarkable ecological diversity, particularly in its occupation of different soil types and various climate zones (*hereafter* referred to as climate niches; Figure 01) [30]. These niches are characterized by large ecological differences involving differences in saltwater exposure, aridity, precipitation, temperature, intense solar radiation. This ecological variation provides an opportunity to explore whether climatic niches exert selective pressures that influence TE dynamics, as observed in plants under arid conditions [14]. Specifically, because the accumulation of TEs can result in an increase in genome size, which is associated with larger stomatal cells with impaired gas exchange efficiency and poor water economy [31], natural selection may regulate genome size in species adapted to arid conditions. Furthermore, hybrid populations exist in the radiation [29], allow testing whether hybridization triggers genome deregulation by inducing TE activity [32–37]. Finally, the reference genome assembly for the group, *Scalesia atractyloides*, is composed of over 80% of repetitive content and TEs – making this lineage ideal to understand TE accumulation dynamics [38].

**Figure 01.**
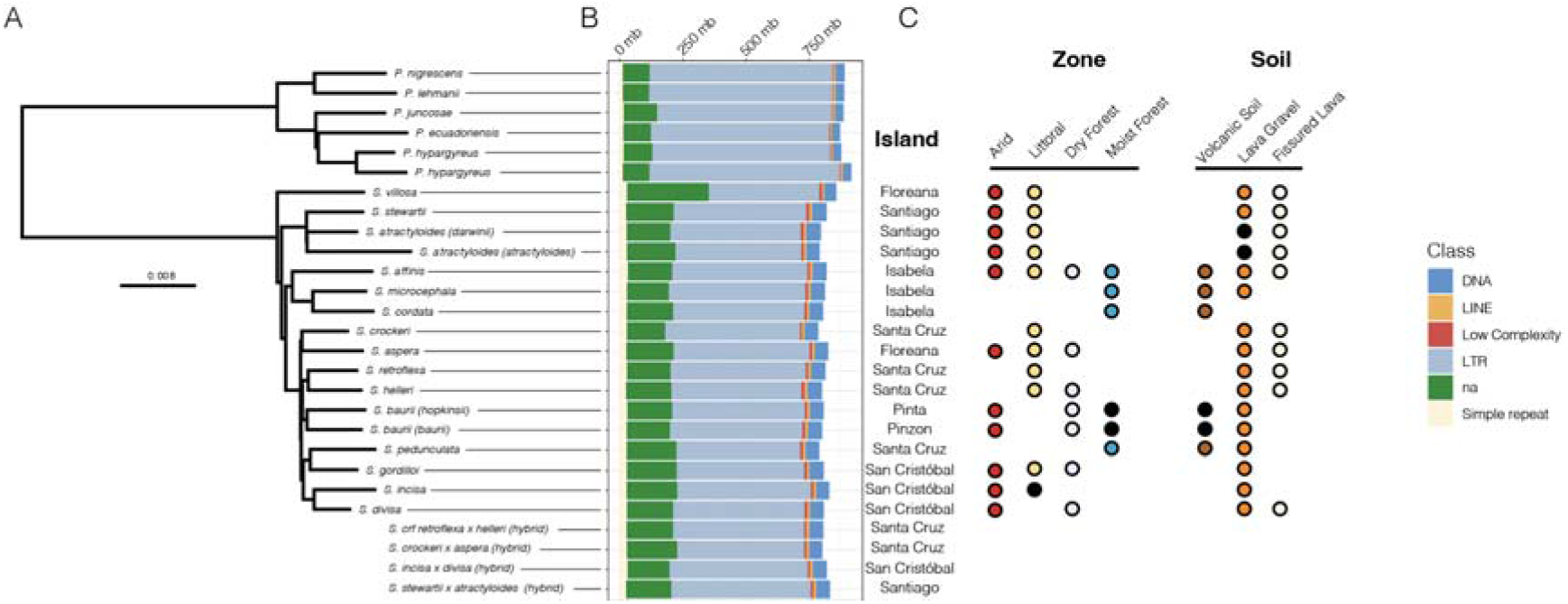
Phylogenetic relationships and transposable element (TE) variation in the *Scalesia* radiation and the outgroup (genus *Pappobolus*). a) Neighbor-joining phylogeny of *Scalesia* and the outgroup. Hybrid samples are shown below and were not included in the phylogenetic reconstruction. b) Stacked bar plot illustrating TE and repeat groups (for the analysis displayed here we used the sunflower library; the analysis using a *Scalesia*-based library is found in Supplementary Figure 02). Color-coding of the different repeat and TE groups is found in the rightmost part of the figure, where ‘na’ denotes unidentified TEs. c) Labelling of the different islands, environments, and soils occupied by different *Scalesia* species (for hybrids we include only island labelling). Environment and soil labelling follow [30], and black dots indicate uncertainty in environmental characterization.

Building on the premise that TEs play a role in diversification dynamics, here we test three hypotheses: *(H1)* Lineages undergoing rapid speciation and broadening of ecological variation exhibit an increased rate of lineage- or taxon-specific TE accumulation/expansion on their genomes; *(H2)* Shifts in climatic niches result in a change of selective regimes, with lineages in arid areas experiencing stronger selection for TE removal as a result of selection for smaller cells; and *(H3)*. Hybrids have an increased accumulation of TEs as a result of genome deregulation. To test these hypotheses, we employed whole-genome resequencing of all 15 *Scalesia* species, four *Scalesia* hybrid populations and outgroup species, and contrasted genomic results with ecological data [30,39].

## Materials and Methods

### Study species and data collection

*Scalesia* (Asteraceae) is an endemic genus to the Galápagos Islands consisting of approximately 15 species derived from a single common ancestor. The divergence of *Scalesia* from members of a continental sister lineage, *Pappobolus*, occurred around 3 million years ago, while most diversification of the group occurred within the last million years [29]. *Scalesia* occupies a wide array of ecological niches and soil types (Figure 1) [30], displaying clear environment-phenotype associations [29,39], as expected in an adaptive radiation [25]. In this study, we sequenced samples representing the entire *Scalesia* genus, including: *S. affinis, S. aspera, S. atractyloides* (var. *atractyloides* and *darwinii*), *S. baurii* (subsp. *baurii* and *hopkinsii*), *S. cordata, S. crockeri, S. divisa, S. gordilloi, S. helleri, S. incisa, S. microcephala, S. pedunculata, S. retroflexa, S. stewartii*, and *S. villosa* (Supplementary Table 01). Additionally, we carried out sequencing of samples from populations that show evidence of extensive hybridization (Bieker, *in prep*). The hybridizing pairs include only pairs of sister species: *S. retroflexa* × *helleri, S. crockerii* × *aspera, S. incisa* × *divisa*, and *S. stewartii* × *atractyloides* (Supplementary Table 01). We incorporated individuals from the genus *Pappobolus* from the South American continent as the outgroup (sister group) in some analyses [29,30], including *P. ecuadoriensis, P. hypargyreus, P. juncosae, P. lehmanii*, and *P. nigrescens*. Detailed information and accession numbers regarding the sampling of outgroup species can be found in Fernández-Mazuecos et al. (2020).

### Data generation (library preparation and sequencing)

We extracted genomic DNA using the DNeasy 96 Plant Kit (Qiagen) following the manufacturer’s protocol with minor modifications. These modifications consisted of grinding dried leaf tissues in a Qiagen collection microtube with a 3-mm tungsten carbide bead and repeating the membrane washing step with an additional 800 µl of buffer AW2 to remove residual greenish color from the samples. We eluted the DNA from the membranes using 50 µl of buffer AE to obtain a high-concentration DNA yield. The DNA extracts were then sent to the commercial provider Novogene UK for shearing to a mean insert size of 350 bp, double-stranded DNA genomic library preparation, indexing and amplification via PCR, and 150-bp paired-end sequencing on the Illumina NovaSeq 6000 platform. The average sequencing depth was ∼6x. Three samples used in this study were obtained from herbarium specimens and were prepared for genomic sequencing independently. Leaf tissue was disrupted using a Tissue Lyser II (Quiagen), followed by DNA extraction using the DNeasy Plant Mini Kit following the manufacturer’s protocol except for an additional overnight incubation step with 20 µL of proteinase K after cell lysis as described in [40]. DNA was eluted using 62 µL buffer AE. Extracts were then sent to the commercial provider Novogene for double-stranded DNA genomic library preparation, indexing and amplification via PCR, and 150-bp paired-end sequencing on the Illumina NovaSeq 6000 platform. The raw sequencing data generated for this study can be found at the European Nucleotide Archive under accession code PRJEB70770.

### Low-coverage phylogenomics

We started by conducting a distance-based phylogenetic reconstruction to investigate the evolutionary history of the *Scalesia* genus and to establish a backbone for subsequent analyses and interpretations. We performed an additional phylogenetic reconstruction, including hybrid populations to confirm the phylogenetic placement of these hybrids. We started by quality-filtering the raw Illumina data by identifying and removing adapters and low-quality basepairs using *AdapterRemoval v2.3.2* [41]. Subsequently, we used *Skmer v3.2.1*, a pipeline designed to estimate genetic distances between genomes through a *k*-mer-based approach. *Skmer* offers an approach for phylogenetic reconstruction without requiring an assembly and accounts for low-coverage of the samples [42–45]. Running *Skmer* involved running the module ‘skmer reference’ to process the adapter-trimmed sequencing data, followed by ‘skmer subsample’ to create 100 subsamples of the libraries, and ‘skmer correct’ to correct the distance matrices based on these subsample replicates. To construct a distance-based phylogenetic tree we employed *fastme v2.1.5* [46].

### TE library construction

We downloaded the reference genome of the sunflower (*Helianthus annuus*) from its official website (https://www.sunflowergenome.org/; version Ha412HOv2.0-20181130), and used it to generate a sunflower-based TE and repeat library using *Repeat Modeler v2.0.2* [47]. Sunflower was selected as the reference species because it has a high-quality reference genome, and it is equally evolutionary divergent from *Scalesia* and *Pappobolus*. Building a TE and repeat library based on *Scalesia* alone could introduce biases when conducting comparisons with the outgroup taxa (but see below). We used the sunflower-based as the base library for subsequent TE detections using *dnaPipeTE*.

We used *dnaPipeTE* [48] to extract and characterize TEs and repeat sequences. *dnaPipeTE* leverages the RNA-seq assembly pipeline Trinity to extract, quantify and assemble sequences, including TEs and other repeats. While the primary objective of *dnaPipeTE* is to construct representative repeat library, it also detects and quantifies partially assembled TE and repeats (which suffice for estimating the relative abundances on which we rely in this study). Upon the completion of the assembly and collection process, *dnaPipeTE* categorises TEs and repeats into different classes based on their homology with the TE and repeat library. In the established sunflower library, TE classes included DNA, Helitron, Long interspersed nuclear element (LINE), Long terminal reads (LTR), rRNA, and Short interspersed nuclear elements (SINE). Leveraging *RepeatMasker, dnaPipeTE* identifies low-complexity sequences, simple repeats, and satellites, guided by a specific coverage threshold. *dnaPipeTE* has been successfully applied to a diverse array of species, including arthropods, mollusks, and vertebrates (reviewed in [49]), even revealing substantial variation in TE and repeat content at the population-level (e.g. [50]). *dnaPipeTE* is designed to work low-coverage data (<1×), identifying TEs and repeats by assembling contigs with high coverage (indicating a high copy number within the genome) and subsequently using a database to classify these genomic structures. To check whether lower coverages would not influence our results, we simulated reads using ART [51], specifying Illumina data, 150 bp read length, 10× coverage, and an insert size of 500 bp with a standard deviation of 50 bp. We simulated reads for three randomly selected chromosomes (12, 19 and 23) and ran *dnaPipeTE* specifying the size of the chromosome and different coverages: (0.1×, 0.2×, 0.5×, 1×, 2×, 5×), three times for each chromosome. This showed that the major classes of TEs/repeats were quite similar (Supplementary Figure 01). We then ran *dnaPipeTE* on single-end forward reads for each individual separately, downscaling the data to 0.5× coverage, specifying a genome size of 3.2 Gbp [38], one Trinity assembly iteration, and a minimal query percentage of 0.2 [48].

### Exploration of patterns

We extracted three outputs from *dnaPipeTE*: 1) counts of TEs and repeats, which allowed a direct comparison of the number of TEs and repeats accumulated across species; 2) repeat landscape plots from all resequenced samples, which offered insights into the accumulation of TEs through time (to complement this data, we also ran a repeat accumulation analysis in the chromosome-resolved *Scalesia atractyloides* genome assembly [38] using RepeatMasker; and 3) fasta files containing TE and repeat sequences. Using the sequences in a fasta format, we used *OrthoFinder* to group TEs and repeats based on sequence similarity [52,53], and examined the number of TE/repeat orthogroups shared between the outgroup species and *Scalesia* using *UpSet plots* and *Venn diagrams*. To test for differences between groups, we performed a Mann–Whitney U test comparing hybrids *vs* non-hybrids, and humid (*S. cordata, S. microcephala, S. pedunculata*) *vs* arid (*S. villosa, S. stewartii, S. atractyloides, S. crockeri, S. aspera, S. retroflexa, S. helleri, S. gordilloi, S. incisa, S. divisa*).

We considered the possibility that the limited variation in TE and repeat content observed in some of our results was due to the use of a sunflower-based library, which might be too divergent from *Scalesia* and *Pappobolus*. To address this concern, we repeated all the aforementioned analyses using a *Scalesia*-specific library instead of the sunflower library. To obtain the *Scalesia*-specific library, we ran *RepeatModeler* on the *Scalesia atractyloides* genome [38] and ran *DNApipeTE* analyses using the same parameters, and running analyses involving the exploration of TE and repeats). Analyses of the *Scalesia*-specific library found similar patterns to the main results (Supplementary Figure 02).

## Results

The phylogenetic tree is comprised mostly of short internal branches in *Scalesia*, as expected in a rapid radiation. *Scalesia villosa* is positioned as the sister species to all the remaining *Scalesia* species (Figure 01A; Supplementary Figure 03). S. *stewartii* and S. *atractyloides* form a clade which is sister to the remaining 12 species. S. *affinis*, S. *microcephala*, and S. cordata are grouped together as a clade being sister to a clade including nine *Scalesia* species, namely *S. crockeri, S. aspera, S. retroflexa, S. helleri, S. baurii, S. pedunculata, S. gordilloi, S. incisa, and S. divisa* (Figure 01A; Supplementary Figure 03). The phylogenetic analysis including hybrid populations placed all hybrids as sister to one of the parental species (Supplementary Figure 04).

We observed little variation in TE counts among different species, both in relative and absolute terms (Figure 01B; Supplementary Table 02). The outgroup lineages displayed a higher count of TEs and repeats, with an average of 1,382,957,707 bp (max 1,398,040,697 bp, min 1,368,357,206 bp, standard deviation 12,799,060 bp). *Scalesia* species (excluding hybrids) exhibited a lower TE and repeat content, with an average of 1,249,051,887 bp (max 1,264,569,945 bp, min 1,223,889,466 bp, standard deviation 1,1922,527 bp). Hybrid *Scalesia* populations displayed a TE and repeat content averaging 1,253,765,218 bp (max 1,265,750,267 bp, min 1,239,589,542 bp, standard deviation 12,633,031 bp; Figure 1B). Both Mann-Whitney U tests were non-significant (humid *vs* arid: W = 16, *P* = 1; hybrid *vs* non-hybrid; W = 41, *P* = 0.5744)

Long terminal repeat (LTR) retrotransposons were the largest class of TEs found in the data. *Scalesia* species (hybrids excluded) had an average of 518,734,809 bp of their genomes composed of LTRs (max 542,466,614 bp, min 439,269,606 bp, standard deviation 24,787,195 bp). The outgroup species had a mean of 719,315,917 bp of their genomes composed of LTRs (max 755,592,383 bp, min 693,803,469 bp, standard deviation 21,926,958 bp), while hybrid *Scalesia* populations had 532,048,025 bp of LTRs (max 553,999,518 bp, min 504,436,424 bp, standard deviation 23,478,054 bp). In the second largest class of TEs, DNA elements, *Scalesia* had an average of 53,397,161 bp composed of DNA elements (max 56,653,585 bp, min 45,126,916 bp, standard deviation 2,748,119 bp), while hybrid populations had 53,374,532 bp (max 54,154,091 bp, min 51,526,434 bp, standard deviation 1,238,711 bp). In contrast, the outgroup exhibited fewer DNA elements, with an average of 32,145,458 bp (max 34,376,438 bp, min 30,424,080 bp, standard deviation 1,414,881 bp).

### Repeat landscape plots

The repeat landscape plot shows that all *Scalesia* species exhibit a similar pattern of TE and repeat accumulation (Figure 02; Supplementary Figure 05). All outgroup species have a similar accumulation of TEs, which is distinct from that of *Scalesia*. Specifically, there is a younger accumulation of TEs in the outgroup relative to *Scalesia* (Figure 02; Supplementary Figure 05). In all the landscape plots, the TE/repeat accumulation peak of *Scalesia* is at a Kimura substitution level of approximately 5 (Figure 02; Supplementary Figures 06). The repeat landscape plots decomposed in different TE and repeat classes show that the accumulation of TEs and repeats does not differ when comparing the outgroup (Supplementary Figure 07), *Scalesia* (Supplementary Figure 08-10), and *Scalesia* hybrids (Supplementary Figure 11).

**Figure 02.**
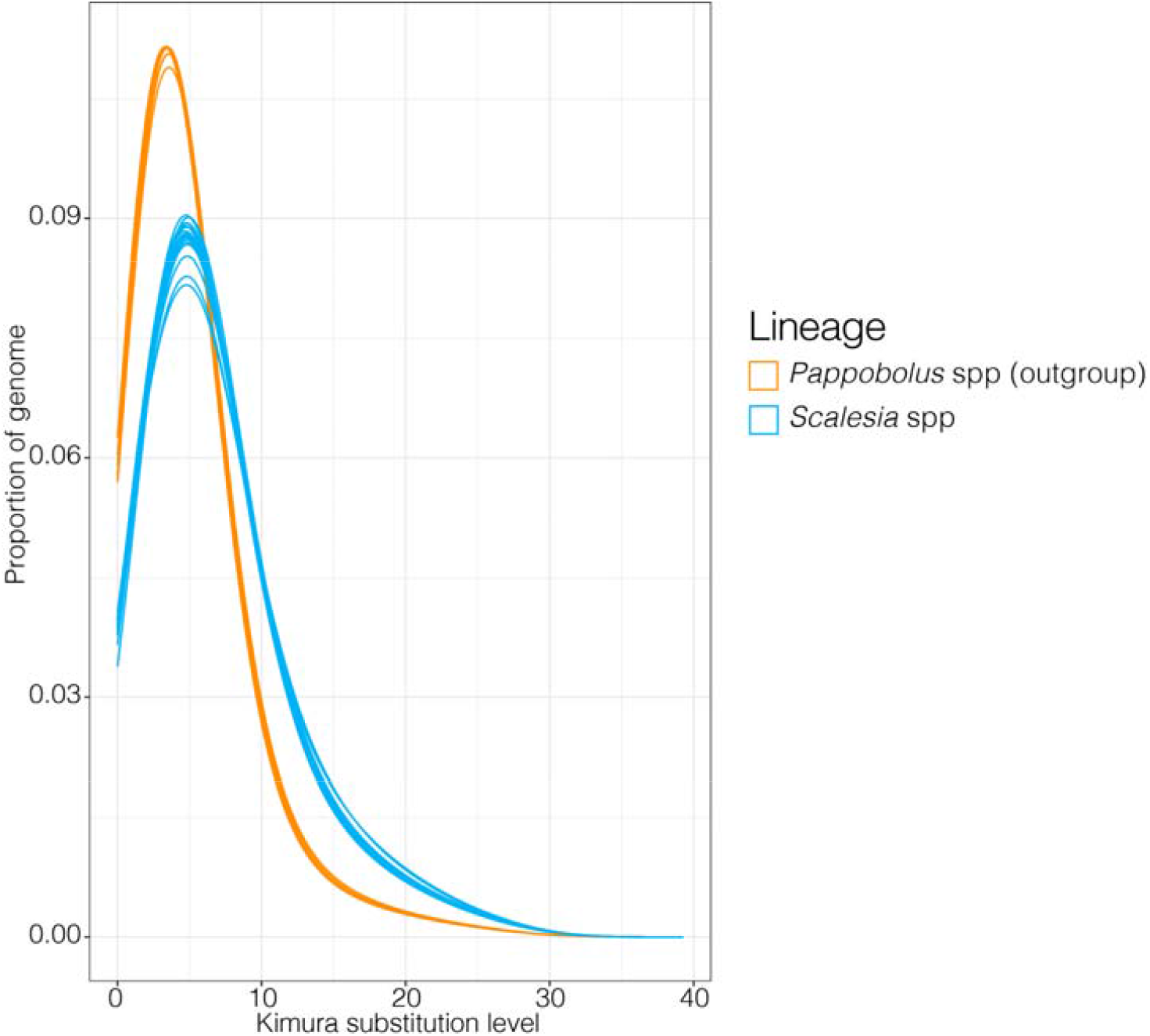
Repeat/transposable element landscape plot. The *x-axis* shows the sequence divergence (Kimura substitution level), an indirect measure of TE divergence/age, the *y-axis* shows the portion of the genome covered by TEs and repeats. Assuming a constant mutation rate, higher Kimura substitution levels indicate older TE ages. The outgroup (genus *Pappobolus*) is plotted in orange, and all *Scalesia* species in blue.

### Organization of genome-specific elements

Using *OrthoFinder*, we grouped TEs and repeats into orthogroups, that are sets of orthologous sequences derived from the last common ancestor of all species under consideration (Figure 02; Supplementary Figures 12-16). In almost all the comparisons, we found that the majority of TE orthogroups are present in all *Scalesia*. The sole exception was in the satellite elements, for which we detected very few shared orthologs. In the largest class, LTR, we found 8,381 orthogroups present in all *Scalesia* genomes, while only 458-255 orthogroups were found to be unique to a single genome (Figure 03).

**Figure 03.**
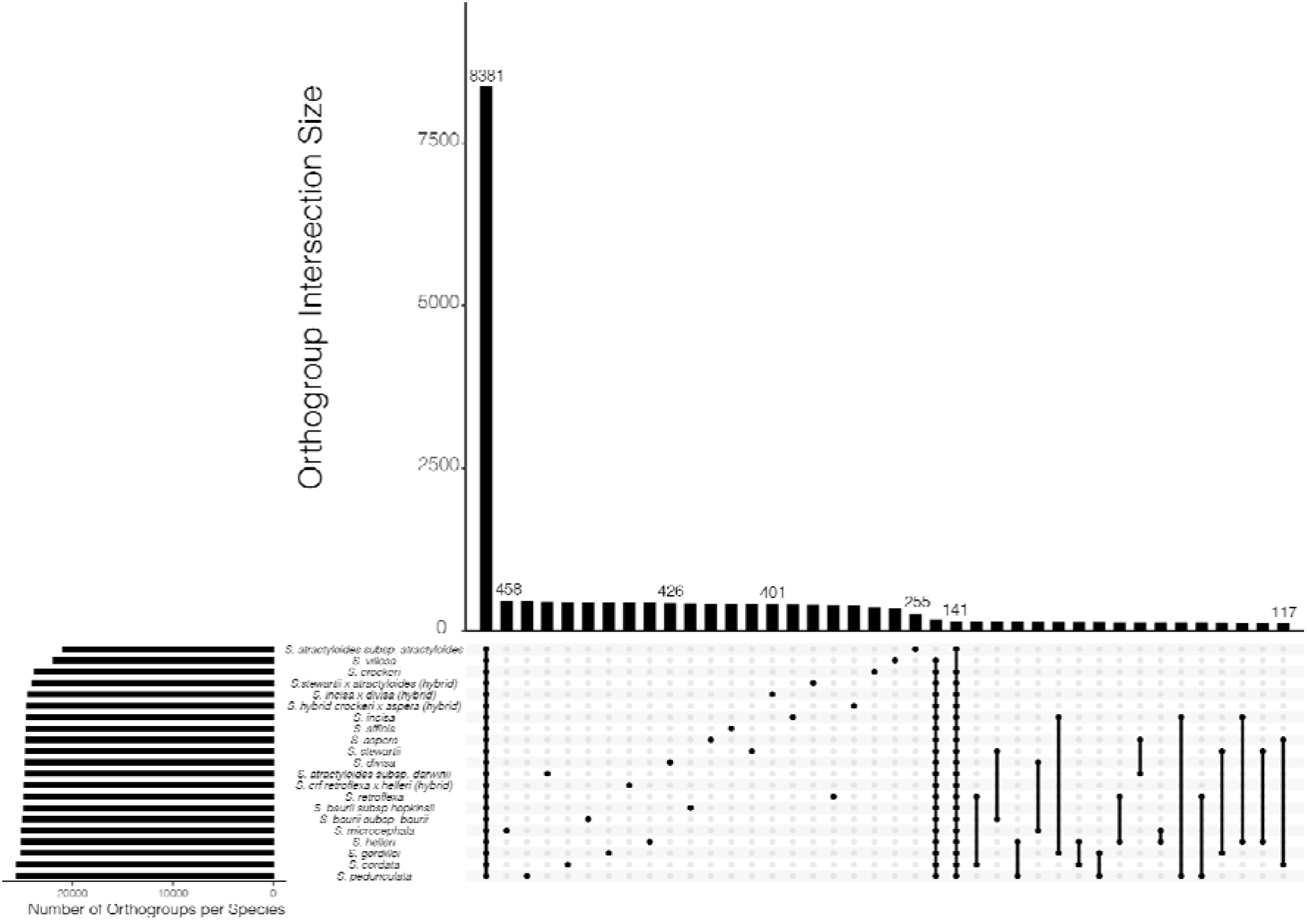
UpSet plot of long terminal repeat elements (LTR) identified within the *Scalesia* radiation (other TE and repeat groups are found in Supplementary Figures 12-16). The *UpSet* plot is an alternative visualisation to a Venn diagram where the y-axis on the top graphic shows a frequency of LTR orthogroups (groups of closely related LTRs) and the x-axis shows different combinations of the data (the first column includes orthogroups present in all *Scalesia*, the second column are orthogroups exclusive to S. *retroflexa x helleri* hybrids, the third column are orthogroups exclusive to S. *atractyloides* var. *atractyloides*). Dark grey circles highlight species combinations.

**[see the end of the document for table 01]**

**Table 01** - TE superfamily counts for different *Scalesia* species.

Superfamily-level analyses show that TE figures were remarkably even in *Scalesia* (Table 01) and the outgroup (Table 01). With the exception of LTR/Gypsy and LTR/Copia, all TE counts were below 50,000 elements and were remarkably even across *Scalesia* species. LTR/Gypsy and LTR/Copia included between 2,500,000 - 983,398 elements (Table 01).

We used *OrthoFinder* to identify orthogroups of TEs and repeats in five randomly chosen *Scalesia* species and five outgroup *Pappobolus* species, and parsed the results to compare counts of *Scalesia*-specific orthogroups, outgroup-specific orthogroups, and orthogroups shared between the two genera (Figure 04). We chose to downscale the data to five random species because we observed imbalances in estimations that arose from analysing the 15 *Scalesia* species in comparison to only five outgroup species (two specimens of P. *hypargyreus* were included;Figure 04). Since we downscaled the data to only five *Scalesia* species, we repeated the analyses three times, finding that the results were consistent across the three iterations (Supplementary Table 03). The largest three classes of TEs (LTR, DNA and Helitron) had a majority of its orthogroups common to *Scalesia* and the outgroup. *Scalesia* had more private SINEs than the outgroup (Figure 04).

**Figure 04.**
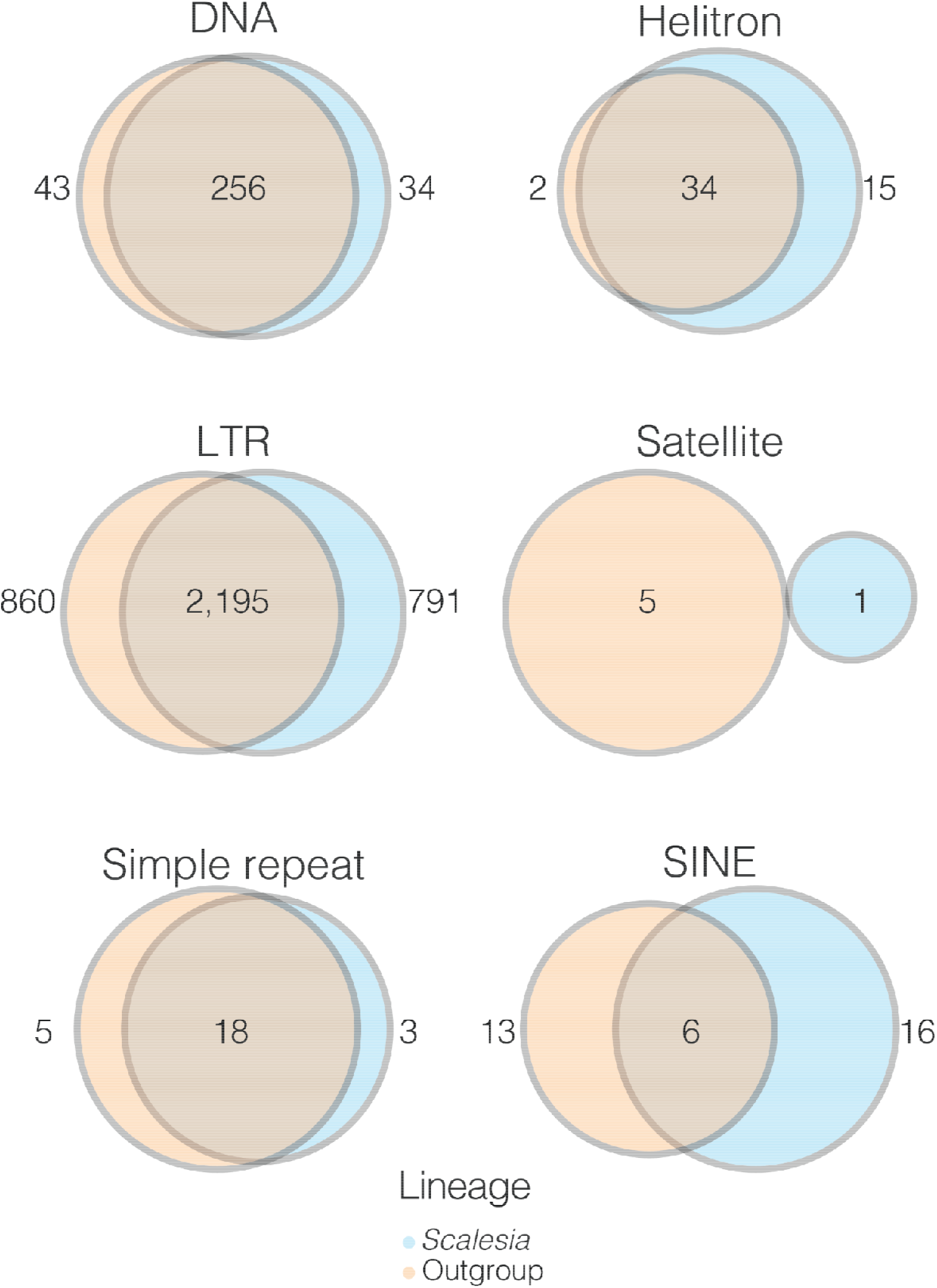
Representation of orthogroups present in *Scalesia* (blue) or the outgroup (orange). Shared orthogroups are given as overlap for different TEs and repeats.

## Discussion

Transposable elements (TEs) and repeats play a crucial role in shaping genomes and phenotype diversity. Here, we tested whether rapid diversification, shifts in climatic niches or hybridization are associated with an increase of TE accumulation in the genomes of the genus *Scalesia*, a radiation endemic to the Galápagos (Figure 01). Our results show that TE or repeat accumulation is consistent across the *Scalesia* phylogeny, with no major changes across species (Figure 01) or over time (Figure 02). We also found no evidence of lineage-specific TE or repeat expansions, as most TE groups were found across all species (Figure 03), in similar numbers across the explored genomes (Table 01), and shared with the outgroup (Figure 04). Overall, our findings indicate that neither rapid diversification, hybridization, nor ecological niche shifts are significantly associated with TE and repeat content variation within the *Scalesia* adaptive radiation.

### Limited variation in transposable element content in Scalesia despite rapid diversification

We found no major differences in TE or repeat content in different *Scalesia* species in any of the analyses performed and focusing on phylogenetic comparisons, accumulation over time, and class-specific expansions (Figures 01-5). The lack of variation may be attributable to the methods used, but this seems unlikely since *dnaPipeTE* retrieved different results for Pappobolus and *Scalesia*, suggesting that the two lineages have accumulated different TEs and repeats after their split, about 3 million years ago [29]. dnaPipeTE has been shown to detect population-level differences [50,54–56], and between closely related species across the Tree of Life [57–63]. One tempting explanation for the lack of variation is that *dnaPipeTE* may be biased to find conserved old elements, thereby explaining the observed patterns. However, this is likely not the case as *dnaPipeTE* is very sensitive in detecting recent TE families, being less sensitive to older TE families due to increased divergence and short reads that limits alignments. For instance, when applied to the human genome, *dnaPipeTE* is biased towards more recent TEs (see supplementary data included in [48]. We therefore suggest that TEs are under phylogenetic inertia, as shown for various *Drosophila* species [64]. Our findings suggest that the fundamental transposable element composition remains conserved across *Scalesia* lineages.

The lack of variation in TE and repeat content in *Scalesia* is noteworthy. In our dataset, we observe a variation of 4.60 % in LTRs, and 4.58 % of variation in DNA elements. This variation is very little when compared to other works focusing on congeneric species. For instance, in the plant genus *Passifora*, Tekay LTR-elements varied between 8.52 - 35.71 % and Angela LTR-elements between 1.96 - 29.05% (absolute estimates considering the whole genome; [65]. In oaks (*Quercus*) LTRs varied between 34.07 - 55.05 % (relative TE content; [66]. In *Rhodnius* bugs, with different ecologies, Tc1/Mariner elements varied between 14.3 - 25 % [63], while in *Diabrotica* beetles MITE elements varied between 150 - 610 MegaBasepairs [62].

One potential explanation for the observed lack of species-specific TEs is the high repeat content of the *Scalesia* genome. Our previous analyses revealed that approximately 80% of the *Scalesia atractyloides* genome is composed of repetitive sequences [38]. This high repeat content could be due to the presence of numerous repeat families that actively replicate with a strong phylogenetic signal, rather than a dynamic turnover of species-specific repeat families with a low copy number. In this regard, Novák and colleagues (2020) found that TE content reaches a plateau at around 80% across angiosperms with genome sizes between 5 and 10 Gbp [67]. It is possible that a plateau of TE content could be maintained, with few families represented at high numbers, and that a dynamic turnover occurs within families, rather than between TE families.

Our findings challenge the notion that lineages with high speciation rates also display elevated rates of TE content accumulation. One caveat to this interpretation is that the sister lineage to *Scalesia*, including *Pappobolus*, may have undergone similar diversification on the mainland [29]. Accordingly, we hypothesize that the entire *Pappobolus-Scalesia* clade possesses ancestrally high TE content. However, recent TE activity is observed as in both lineages most TEs are relatively young (Figure 02). In any case, the role of TEs in adaptive radiation remains debated, with some studies demonstrating their importance as drivers of radiation [68,69], while others proposing a less significant role [27,28]. This discrepancy could stem from the scale and data employed. Specifically, the correlation between TE content and diversification based on limited data may have led to an overestimation of TEs’ role. For instance, an initial analysis of four transcriptomes and genomes of East African cichlids, compared to tilapia and other teleosts, suggested a higher accumulation of TE counts during the radiation [26]. In contrast, recent evidence involving analyses of hundreds of genomes found no relationship between species diversity and TEs within different tribes of the Tanganyika radiation [27]. However, this does not negate the compelling experimental evidence supporting the role of TEs in driving the evolution of specific phenotypic change, such as the elegant findings of Kratochwil and colleagues demonstrating that a TE insertion triggered the emergence of a novel phenotype, the cichlid gold morph [11]. Considering this, we suggest that caution should be taken with potential overestimation of the role of TEs as major driving forces in speciation and ecological diversification at the macroevolutionary scale in all cases (e.g. [15–22,70–72]). This is particularly evident given the limitations of existing studies, which often rely on limited genomic data, correlational analyses, and a narrow taxonomic scope. As a result, some of these correlations encompassing macroevolutionary scales might have overstated their impact on adaptive evolution, speciation, and their direct connection to evolutionary success. Indeed, it can be argued that TEs are unlikely to enhance diversification unless genetic variation is limiting, such as in a bottleneck – as expected in lineages such as *Scalesia*. While TEs offer a convenient explanation for rapid evolution, it is likely that they are merely “along for the ride” and may only be purged when they reach problematic levels of replication.

### TEs and ecological variation

The minimal variation in TE accumulation observed in *Scalesia* suggests that the climatic niche does not significantly influence TE content within the radiation. In adaptive radiations, closely related species typically occupy distinct ecological niches [24,25,73]. In *Scalesia*, species exhibit distinct climatic niches (Figure 01) [29,30,39], with temperature and the presence of arid environments being key factors shaping the evolution of the radiation [29,38,39]. Various studies have demonstrated that aridity exerts a significant selective pressure on plant genome size reduction (e.g. [14,74], as this correlates nearly 1-to-1 with smaller cell sizes [75]. Smaller cells improve water economy as smaller stomata respond more efficiently to water stress, enhance gas exchange regulation, and optimize photosynthetic performance [31,76,77]. Given that arid environments favor smaller cells, aridity is hypothesized to be an important selective pressure regulating genome size [14]. Therefore, it would be expected that TEs are being purged in species occupying arid coastal areas (e. g. S. *affinis*), where temperatures frequently reach 40°C throughout the year and water stress is pervasive, compared to species found in the humid highlands (e.g., S. *pedunculata*), where conditions are consistently moist and cooler. This leads to an alternative hypothesis: perhaps selection is unable to effectively remove transposable elements. The removal of a single TE, only a few kilobases in size, might be ‘invisible’ in a genome of 3-4 gigabases [78].

### TEs and hybridization

The four hybrids examined in this study exhibited similar TE content to non-hybrid populations, excluding the possibility of genome deregulation through TE-release as a result of hybridization. Traditionally, hybridization has been perceived as a detrimental force to the integrity of species [79], and it has been proposed that hybridization may lead to genome destabilisation by promoting TE activity [1,80]. The presence of more TEs in hybrid populations would be expected in a scenario involving genome shock triggered by multi-generational hybridization. In such cases, increased TE transcription should lead to more TEs insertions in the hybrid genomes, and a consequent increase of genome size. However, the case in *Scalesia*, as evidenced by the absence of supporting patterns in the repeat accumulation curves (Figure 02), the lack of superfamily expansion or super-family level (Figure 03), and absolute accumulation (Figure 01). These findings are in agreement with those of [81], who demonstrated no evidence of LTR retrotransposon proliferation when comparing multiple hybrid populations of *Helianthus annuus* (sunflower) × *H. petiolaris*. This result suggests that post-transcriptional repression mechanisms may prevent the proliferation of LTRs [81]. A similar scenario may be at play in *Scalesia*, but additional data would be required to confirm this hypothesis.

## Acknowledgements

We were supported by the Research Council of Norway, project 287327. We used the supercomputer Saga, managed by Sigma2 (National e-infrastructure services of Norway; projects nn10082k, and nn9449k). Samples fromf *Scalesia* populations were originally obtained in accordance with Galápagos National Park research permit number PC-001/98 PNG, and later subjected to normalisation procedures authorized by the Ecuadorian Ministry of the Environment genetic permit number MAAE-DBI-CM-2021-0213. This publication is contribution number 2613 of the Charles Darwin Foundation for the Galápagos Islands. We are grateful to Alexander Suh, William B. Reinar and Irene Julca for support in TE analyses and for their helpful comments. We are also thankful to Ross McCauley for providing the *Pappobolus* samples and to Justine Villalba for assisting with the sampling of the *Scalesia* herbarium specimens.

