## Supplementary for "Genomic stability in the Galápagos *Scalesia* adaptive radiation: Consistent transposable element accumulation despite hybridization and ecological niche shifts" (from another version of this preprint: version 1, 2024-10-01)

**Supplementary Table 01** Specimens used with sampling coordinates and information

| **Species** | **Individual identifier** | **Sampling location** | **Island** | **Collection date** |
| --- | --- | --- | --- | --- |
| *Scalesia villosa* | HAMP 24_22 | 1° 13' 55.0"S  90° 21' 41.0"W | Floreana | 1998-01-20 |
| *S. stewartii* | HALM 36_6 | 0° 17' 21.9'' S  90° 34' 27.4'' W | Santiago | 1999-02-03 |
| *S. atractyloides subs. darwinii* | CDRS 17604 | 0° 17' 52.8966" S 90° 38' 31.6968" W | Santiago | 2005-07-18 |
| *S. atractyloides subsp. atractyloides* | CDRS 16393 | 0° 18' 27.8958" S 90° 37' 17.9976" W | Santiago | 2004-02-26 |
| *S. affinis* | HALM 48_21 | 0° 55' 18.3''S  90° 59' 18.5''W | Isabela | 1999-02-11 |
| *S. microcephala* | HALM 51_23 | (Darwin Volcano) | Isabela | 1999-02-06 |
| *S. cordata* | HALM 44_18 | 0° 50' 48.1'' S  91° 01' 18.9'' W | Isabela | 1999-02-10 |
| *S. crockeri* | HALM 39_23 | 0° 34' 45.1'' S  90° 10' 22.8'' W | Santa Cruz | 1999-02-06 |
| *S. aspera* | HALM 37_13 | 0° 33' 36.3'' S  90° 31' 59.9'' W | Floreana | 1999-02-03 |
| *S. retroflexa* | HALM 26_22 | 0° 44' 40.7'' S  90° 16' 22.1'' W | Santa Cruz | 1999-01-26 |
| *S. helleri* | HALM 62_14 | 0° 45' 15.5'' S  90° 25' 26.8'' W | Santa Cruz | 1999-05-24 |
| *S. baurii subsp. hopkinsii* | SLAM 20_17 | 0° 31' 51.1'' N  90° 44' 19.6'' W | Pinta | 2000-01-11 |
| *S. baurii subsp. baurii* | AHA 6_15 | 0° 36' 23'' S  90° 40' 12'' W | Pinzon | 2004-02-16 |
| *S. pedunculata* | HALM 25_1 | 0° 37' 27'' S  90° 23' 02'' W | Santa Cruz | 1999-01-25 |
| *S. gordilloi* | HAMP 38_18 | 0° 55' 49'' S  89° 36' 29'' W | San Cristóbal | 1998-01-26 |
| *S. incisa* | HALM 20_28 | 0° 41' 44.4'' S  89° 18' 03.4'' W | San Cristóbal | 1999-01-21 |
| *S. divisa* | HALM 11_8 | 0° 47' 26.2'' S  89° 20' 07.7'' W | San Cristóbal | 1999-01-17 |
| *S. retroflexa x helleri (hybrid)* | HALM 41_22 | 0° 41' 40.1'' S  90° 11' 28.2'' W | Santa Cruz | 1999-02-06 |
| *S. crockeri x aspera (hybrid)* | HALM 8_22 | 0° 32' 21.2'' S  90° 19' 01.6'' W | Santa Cruz | 1999-01-15 |
| *S. incisa x divisa (hybrid)* | HALM 23_26 | 0° 49' 07'' S 89° 21’ 12’’ W | San Cristóbal | 1999-01-13 |
| *S. stewartii x atractyloides (hybrid)* | CDRS 17571 | - | - | - |

**Supplementary Table 02** The total number of base pairs encompassed within the genomes of various species by distinct TEs and repeats, as determined using DNApipeTE. Megabasepairs (Mb) are given in parenthesis.

| **Species** | **DNA** | **Helitron** | **LINE** | **Low Complexity** | **LTR** | **na** | **rRNA** | **Satellite** | **Simple repeat** | **SINE** | **others** |
| --- | --- | --- | --- | --- | --- | --- | --- | --- | --- | --- | --- |
| *Scalesia villosa* | 11743596  (11,7 Mb) | 888806  (0.9 Mb) | 2514436  (2.5 Mb) | 662977  (0.7 Mb) | 106894927  (106.9 Mb) | 59404313  (59.4 Mb) | 1592119  (1.6 Mb) | 196558  (0.2 Mb) | 3713043  (3.7 Mb) | 1350714  (1.4 Mb) | 80332390  (80.3 Mb) |
| *S. stewartii* | 13464511  (13,5 Mb) | 645022  (0.6 Mb) | 3082091  (3.1 Mb) | 1133364  (1.1 Mb) | 127485403  (127,4 Mb) | 24742002  (24.7 Mb) | 1422234  (1.4 Mb) | 692334  (0.7 Mb) | 3983945  (4 Mb) | 1520775  (1.5 Mb) | 90753318  (90.7 Mb) |
| *S. atractyloides subs. darwinii* | 14284872  (14,3 Mb) | 555759  (0.6 Mb) | 3532521  (3.5 Mb) | 1198062  (1.2 Mb) | 126868393  (126,9 Mb) | 22968899  (23 Mb) | 1016582  (1 Mb) | 419140  (0.4 Mb) | 3696825  (3.7 Mb) | 1792256  (1.8 Mb) | 92932261  (93 Mb) |
| *S. atractyloides subsp. atractyloides* | 11797972  (11,8 Mb) | 510364  (0.5 Mb) | 2956690  (3 Mb) | 1029776  (1 Mb) | 122932407  (123 Mb) | 32782585  (32.8 Mb) | 1422242  (1.4 Mb) | 375175  (0.4 Mb) | 4553943  (4.5 Mb) | 1457081  (1.5 Mb) | 90026739  (90 Mb) |
| *S. affinis* | 13515053  (13,5 Mb) | 756331  (0.8 Mb) | 3327448  (3.3 Mb) | 839702  (0.8 Mb) | 130687970  (130,1 Mb) | 21941234  (22 Mb) | 1290066  (1.3 Mb) | 794659  (0.8 Mb) | 3440621  (3.4 Mb) | 1651739  (1.7 Mb) | 91751804  (91.8 Mb) |
| *S. microcephala* | 13629529  (13,6 Mb) | 735336  (0.7 Mb) | 2742483  (2.7 Mb) | 897553  (0.9 Mb) | 130208566  (130,2 Mb) | 18801796  (18.8 Mb) | 2175086  (2.2 Mb) | 507955  (0.5 Mb) | 3289418  (3.3 Mb) | 1488020  (1.5 Mb) | 93838664  (93.8 Mb) |
| *S. cordata* | 13121661  (13,1 Mb) | 620445  (0.6 Mb) | 2866000  (2.9 Mb) | 1076758  (1.1 Mb) | 124842930  (124,8 Mb) | 22497357  (22.5 Mb) | 819872  (0.8 Mb) | 667639  (0.7 Mb) | 3738872  (3.7 Mb) | 1147854  (1.1 Mb) | 92782812  (92.8 Mb) |
| *S. crockeri* | 13379779  (13,4 Mb) | 838464  (0.8 Mb) | 2952952  (3 Mb) | 1070746  (1.1 Mb) | 130280488  (130,3 Mb) | 17991625  (18 Mb) | 1594736  (1.6 Mb) | 416244  (0.4 Mb) | 3695577  (3.7 Mb) | 1694337  (1.7 Mb) | 91363252  (91.4 Mb) |
| *S. aspera* | 13544776  (13,5 Mb) | 766621  (0.8 Mb) | 3370161  (3.4 Mb) | 963577  (1 Mb) | 129606870  (129,6 Mb) | 25688690  (25.7 Mb) | 2030075  (2 Mb) | 624705  (0.6 Mb) | 3376624  (3.4 Mb) | 1615455  (1.6 Mb) | 90356274  (90.4 Mb) |
| *S. retroflexa* | 12954550  (13 Mb) | 745498  (0.7 Mb) | 3850825  (3.9 Mb) | 604710  (0.6 Mb) | 129804663  (129,8 Mb) | 20511455  (20.5 Mb) | 995881  (1 Mb) | 418176  (0.4 Mb) | 3713353  (3.7 Mb) | 1456998  (1.5 Mb) | 93726444  (93.7 Mb) |
| *S. helleri* | 13115364  (13,1 Mb) | 694038  (0.7 Mb) | 3378061  (3.4 Mb) | 1663852  (1.6 Mb) | 125843322  (126,8 Mb) | 19447114  (19.5 Mb) | 1038917  (1 Mb) | 396329  (0.4 Mb) | 3657768  (3.7 Mb) | 1377096  (1.4 Mb) | 90102529  (90.1 Mb) |
| *S. baurii subsp. hopkinsii* | 14342757  (14,3 Mb) | 705997  (0.7 Mb) | 2720020  (2.7 Mb) | 1518811  (1.5 Mb) | 127558529  (127,6 Mb) | 20720854  (20.7 Mb) | 1475428  (1.5 Mb) | 467590  (0.5 Mb) | 3944576  (4 Mb) | 1173055  (1.2 Mb) | 91573370  (91.6 Mb) |
| *S. baurii subsp. baurii* | 13334489  (13,3 Mb) | 539181  (0.5 Mb) | 2714358  (2.7 Mb) | 1085489  (1.1 Mb) | 127659575  (127,7 Mb) | 19612786  (19.6 Mb) | 1479226  (1.5 Mb) | 786922  (0.8 Mb) | 3889583  (3.9 Mb) | 1163806  (1.2 Mb) | 94236425  (94.2 Mb) |
| *S. pedunculata* | 13279313  (13,3 Mb) | 610044  (0.6 Mb) | 2610080  (2.6 Mb) | 828382  (0.8 Mb) | 117633237  (117,6 Mb) | 24623061  (24.6 Mb) | 1075217  (1.1 Mb) | 347253  (0.3 Mb) | 3655296  (3.7 Mb) | 1630564  (1.6 Mb) | 93405752  (93.4 Mb) |
| *S. gordilloi* | 12312598  (12,3 Mb) | 679900  (0.7 Mb) | 2591697  (2.6 Mb) | 1082070  (1.1 Mb) | 123228827  (123,2 Mb) | 25115012  (25.1 Mb) | 979113  (1 Mb) | 409068  (0.4 Mb) | 4407217  (4.4 Mb) | 1373448  (1.4 Mb) | 89355297  (89.4 Mb) |
| *S. incisa* | 12974348  (13 Mb) | 713527  (0.7 Mb) | 2607612  (2.6 Mb) | 1075471  (1.1 Mb) | 127928243  (127,9 Mb) | 26704393  (26.7 Mb) | 1490230  (1.5 Mb) | 544678  (0.5 Mb) | 3582431  (3.6 Mb) | 1360585  (1.4 Mb) | 91699378  (91.7 Mb) |
| *S. divisa* | 14561033  (14,6 Mb) | 695160  (0.7 Mb) | 3012856  (3 Mb) | 1079731  (1.1 Mb) | 124567696  (124,6 Mb) | 22739517  (22.8 Mb) | 886468  (0.9 Mb) | 635378  (0.6 Mb) | 4043408  (4 Mb) | 1726820  (1.7 Mb) | 92799082  (91.8 Mb) |
| *S. retroflexa x helleri (hybrid)* | 13801838  (13,8 Mb) | 589648  (0.6 Mb) | 3380374  (3.4 Mb) | 834857  (0.8 Mb) | 125159196  (125,2 Mb) | 23688830  (23.7 Mb) | 1109171  (1.1 Mb) | 301222  (0.3 Mb) | 3826935  (3.8 Mb) | 1658986  (1.7 Mb) | 91078298  (91.1 Mb) |
| *S. crockeri x aspera (hybrid)* | 12819925  (12,8 Mb) | 626331  (0.6 Mb) | 2879781  (2.9 Mb) | 1105887  (1.1 Mb) | 122925507  ( 123 Mb) | 28531136  (28.5 Mb) | 1753624  (1.8 Mb) | 454933  (0.5 Mb) | 3517273  (3.5 Mb) | 1602564  (1.6 Mb) | 89667395  (89.7 Mb) |
| *S. incisa x divisa (hybrid)* | 14377105  (14,4 Mb) | 1006770  (1 Mb) | 3421054  (3.4 Mb) | 793055  (0.8 Mb) | 134761987  (134,8 Mb) | 21425773  (21.4 Mb) | 1573311  (1.6 Mb) | 523349  (0.5 Mb) | 3578604  (3.6 Mb) | 1319441  (1.3 Mb) | 90615040  (90.6 Mb) |
| *S. stewartii x atractyloides (hybrid)* | 12542486  (12,5 Mb) | 593503  (0.6 Mb) | 3546128  (3.5 Mb) | 894861  (0.9 Mb) | 135389443  (135,4 Mb) | 24744557  (24.8 Mb) | 1813426  (1.8 Mb) | 712058  (0.7 Mb) | 4081581  (4.1 Mb) | 1401138  (1.4 Mb) | 88735103  (88.7 Mb) |
| *P. ecuadoriensis (outgroup)* | 7308486  (7,3 Mb) | 908968  (0.9 Mb) | 2256582  (2.2 Mb) | 419838  (0.4 Mb) | 172328395  (172,3 Mb) | 20797599  (20.8 Mb) | 2016348  (2 Mb) | 301727  (0.3 Mb) | 2489975  (2.5 Mb) | 139148  (0.1 Mb) | 112160209  (112.2 Mb) |
| *P. hypargyreus (outgroup)* | 7465784  (7,5 Mb) | 954690  (1 Mb) | 2857293  (2.9 Mb) | 168451  (0.1 Mb) | 178110682  (178,1 Mb) | 18868469  (18.9 Mb) | 2391825  (2.4 Mb) | 874972  (0.9 Mb) | 2179691  (2.2 Mb) | 126584  (0.1 Mb) | 111887727  (111.9 Mb) |
| *P. hypargyreus (outgroup)* | 7598123  (7,6 Mb) | 1026287  (1 Mb) | 2739217  (2.7 Mb) | 287744  (0.3 Mb) | 170325380  (170,3 Mb) | 22104009  (22.1 Mb) | 1568471  (1.6 Mb) | 253863  (0.3 Mb) | 2462565  (2.5 Mb) | 132609  (0.1 Mb) | 111948829  (112 Mb) |
| *P. juncosae (outgroup)* | 7625127  (7,6 Mb) | 1124446  (1.1 Mb) | 2288246  (2.3 Mb) | 249888  (0.2 Mb) | 166411171  (166,4 Mb) | 24819745  (24.9 Mb) | 3642726  (3.6 Mb) | 436540  (0.4 Mb) | 2838295  (2.8 Mb) | 143382  (0.1 Mb) | 108900718  (109 Mb) |
| *P. lehmanii (outgroup)* | 7967219  (8 Mb) | 1015156  (1 Mb) | 2523170  (2.5 Mb) | 263764  (0.3 Mb) | 172851793  (172,9 Mb) | 20203839  (20.2 Mb) | 1737311  (1.7 Mb) | 249697  (0.2 Mb) | 2108039  (2.1 Mb) | 119476  (0.1 Mb) | 119121456  (119.1 Mb) |
| *P. nigrescens (outgroup)* | 8258881  (8,3 Mb) | 1152523  (1.2 Mb) | 2355336  (2.3 Mb) | 258758  (0.3 Mb) | 177153648  (177,2 Mb) | 19695728  (19.7 Mb) | 2625878  (2.6 Mb) | 486176  (0.5 Mb) | 2327311  (2.3 Mb) | 140221  (0.1 Mb) | 110849904  (110.9 Mb) |

**Supplementary Table 03** We ran OrthoFinder analyses on 5 random *Scalesia* and 5 random outgroup genomes three times to assess the consistency of our results. Run 1 is displayed in Figure 3.

| **Lineage** | **DNA** | **Helitron** | **LTR** | **SINE** | **Satelite** | **Simple repeat** |
| --- | --- | --- | --- | --- | --- | --- |
| **Run 1** |  |  |  |  |  |  |
| *Scalesia* | 18 | 4 | 464 | 12 | 3 | 1 |
| *Outgroup* | 13 | 5 | 521 | 1 | 5 | 0 |
| *Common* | 99 | 13 | 1067 | 2 | 2 | 1 |
| **Run 2** |  |  |  |  |  |  |
| *Scalesia* | 18 | 4 | 504 | 12 | 1 | 2 |
| *Outgroup* | 11 | 6 | 511 | 1 | 1 | 1 |
| *Common* | 110 | 8 | 1094 | 5 | 1 | 2 |
| **Run 3** |  |  |  |  |  |  |
| *Scalesia* | 19 | 3 | 456 | 15 | 2 | 2 |
| *Outgroup* | 13 | 5 | 528 | 2 | 5 | 0 |
| *Common* | 96 | 13 | 1081 | 2 | 1 | 5 |

**Supplementary Figure 01** Stacked bar plot illustrating various TE classes (legend provided in the right part of the corner, “na*”* denotes unidentified TEs), for the analysis done with the *Scalesia-*specific TE library.

**Supplementary Figure 02 Supplementary Figure 02** Neighbor-joining phylogenetic tree of *Scalesia* radiation and the outgroup. This tree is the same as the tree presented as part of Figure 01 in the main paper, but here specimen identifiers are included.

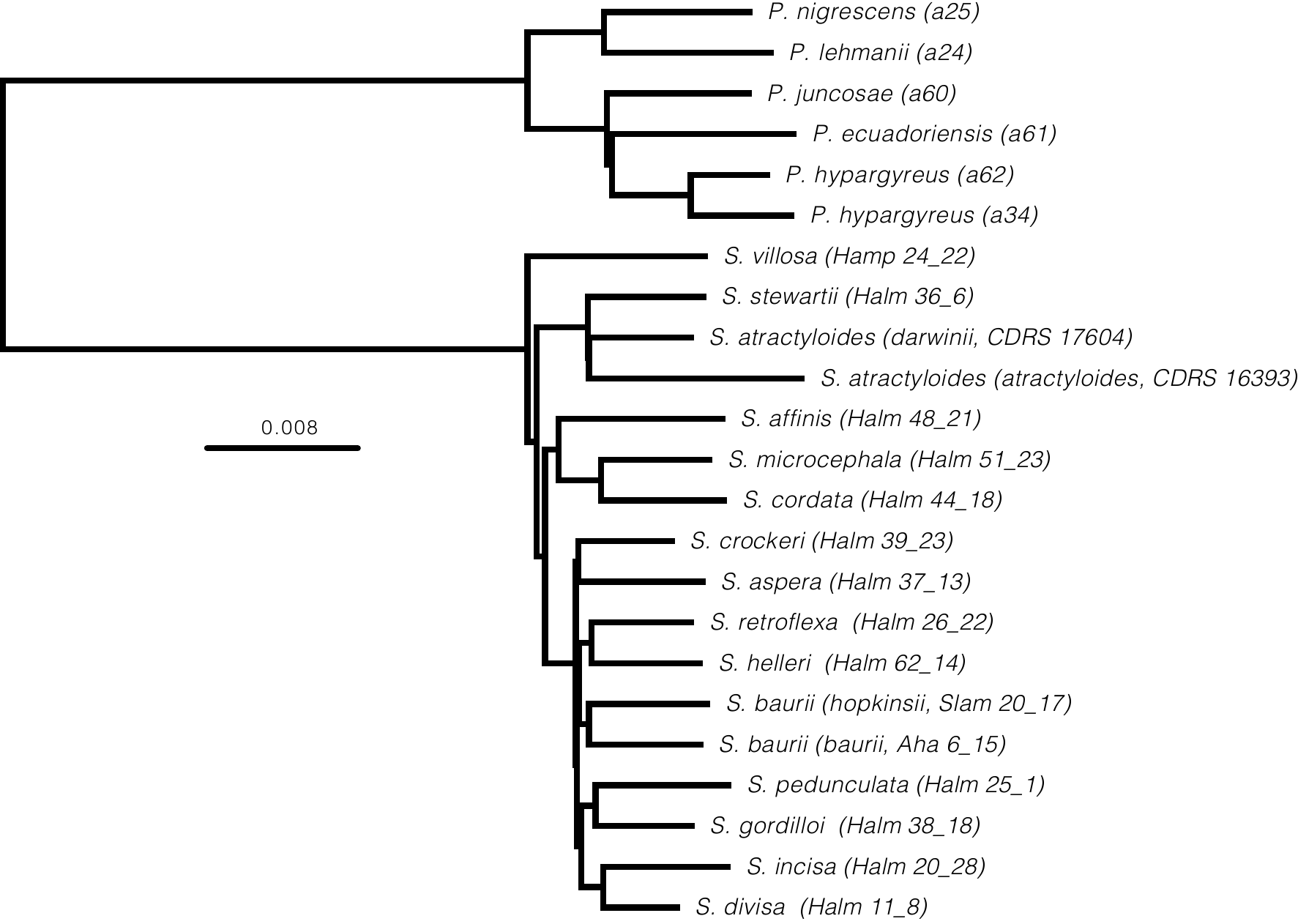

**Supplementary Figure 03** Neighbor-joining phylogenetic tree of *Scalesia* radiation and the outgroup. This analysis includes hybrid specimens (bold).

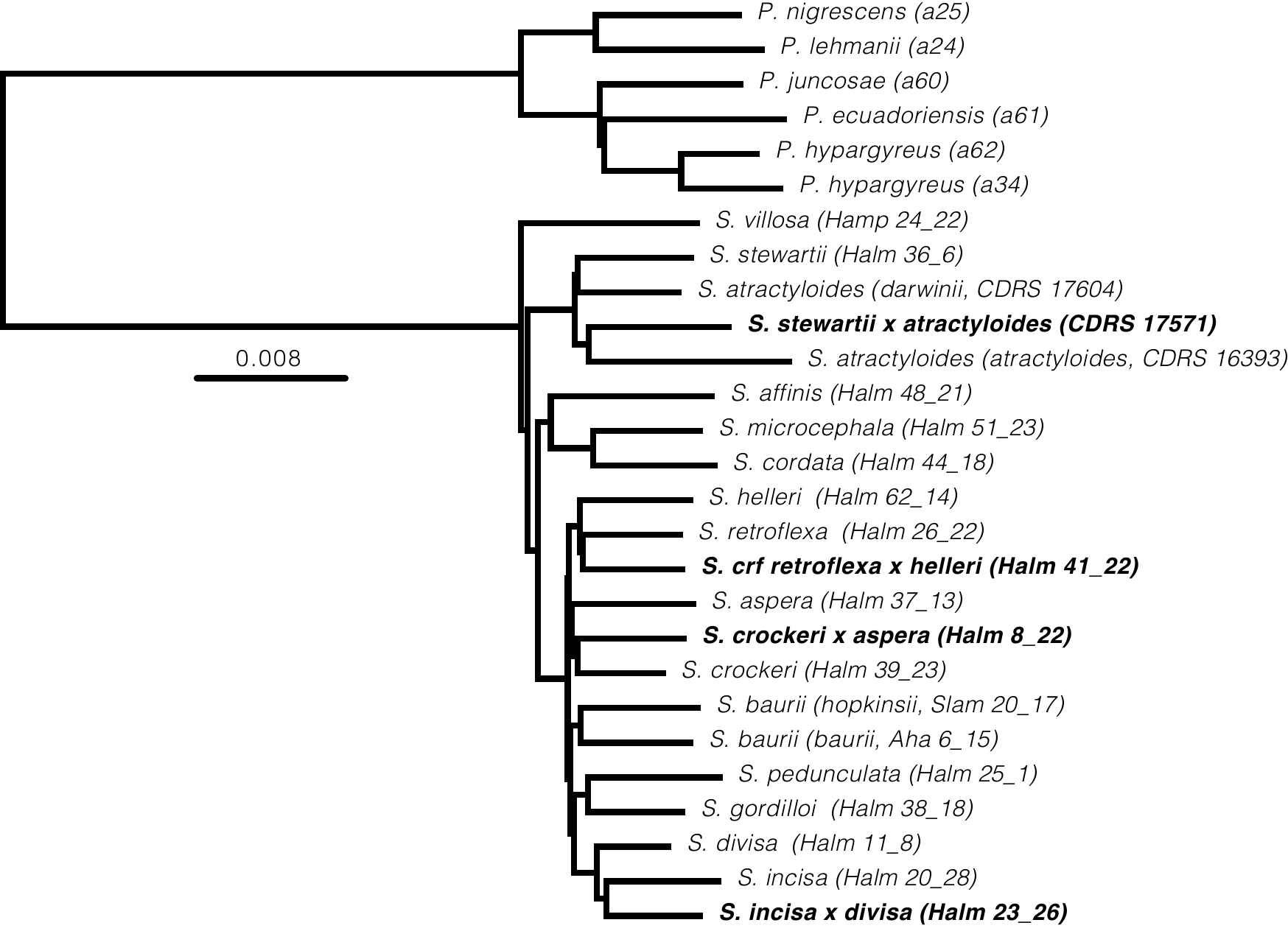

**Supplementary Figure 04** Repeat landscape plot showing the divergence (Kimura substitution-level) of transposable elements (TEs) in the genome. This plot is similar to Figure 02, but each species is represented by a different colour.

**Supplementary Figure 05** Repeat landscape plot of the *Scalesia atractyloides* reference genome.
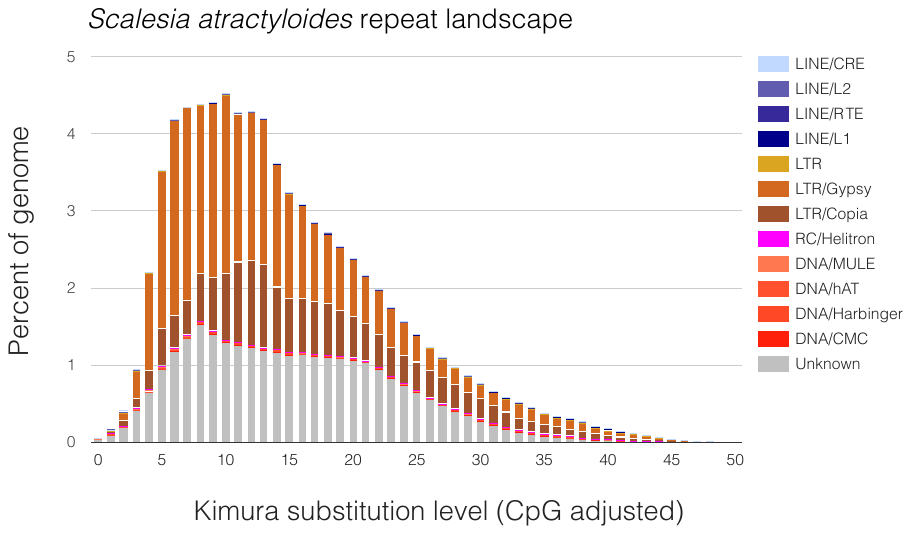

**Supplementary Figure 06** Repeat landscape plot showing the divergence of transposable elements (TEs) in the genome for each of the outgroup specimens. Different TE groups are represented by different colors.

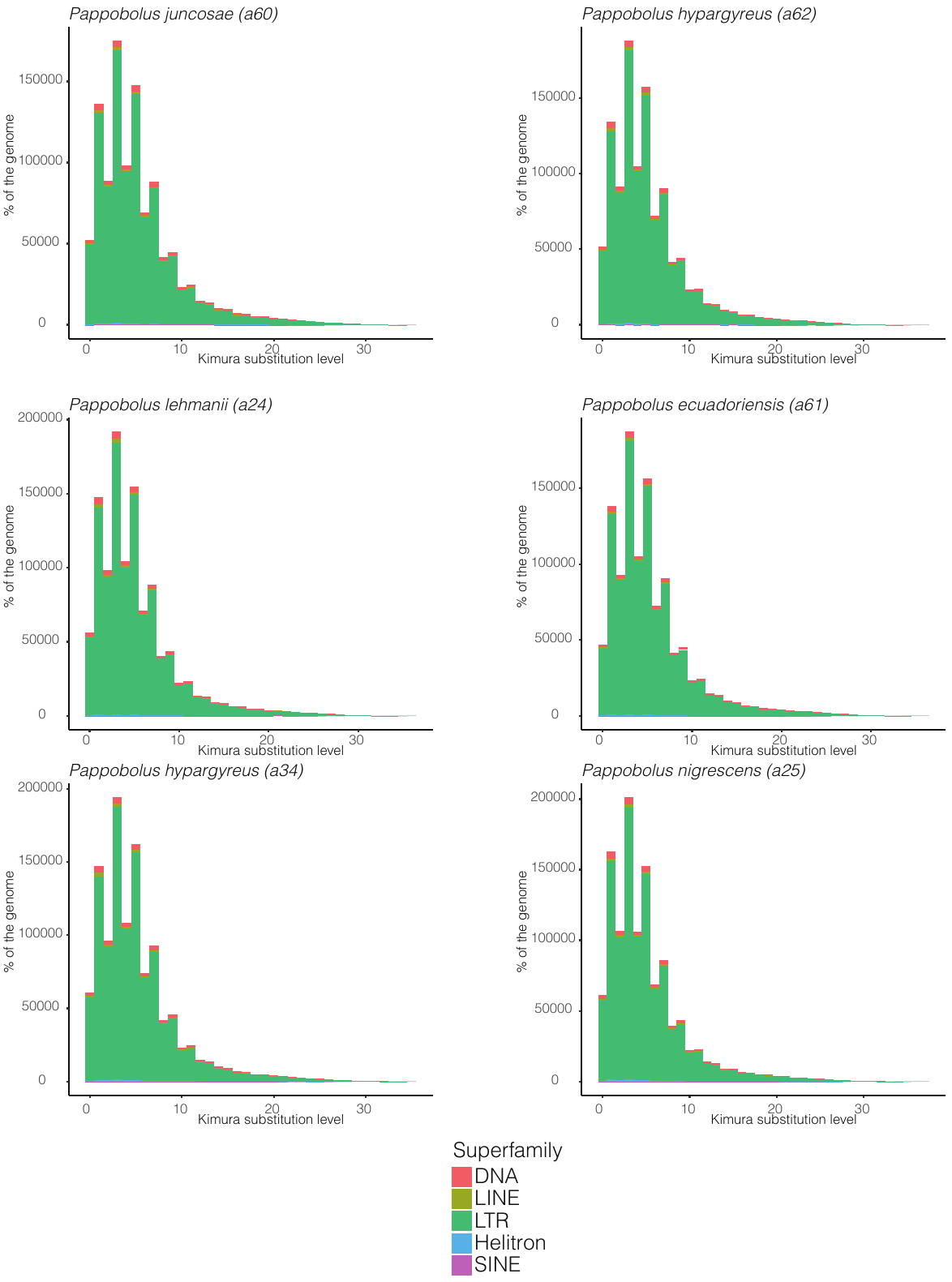

**Supplementary Figure 07** Repeat landscape plot showing the divergence of transposable elements (TEs) in the genome for *Scalesia* *affinis, S. aspera, S. atractyloides and S. baurii*. Different TE groups are represented by different colors.

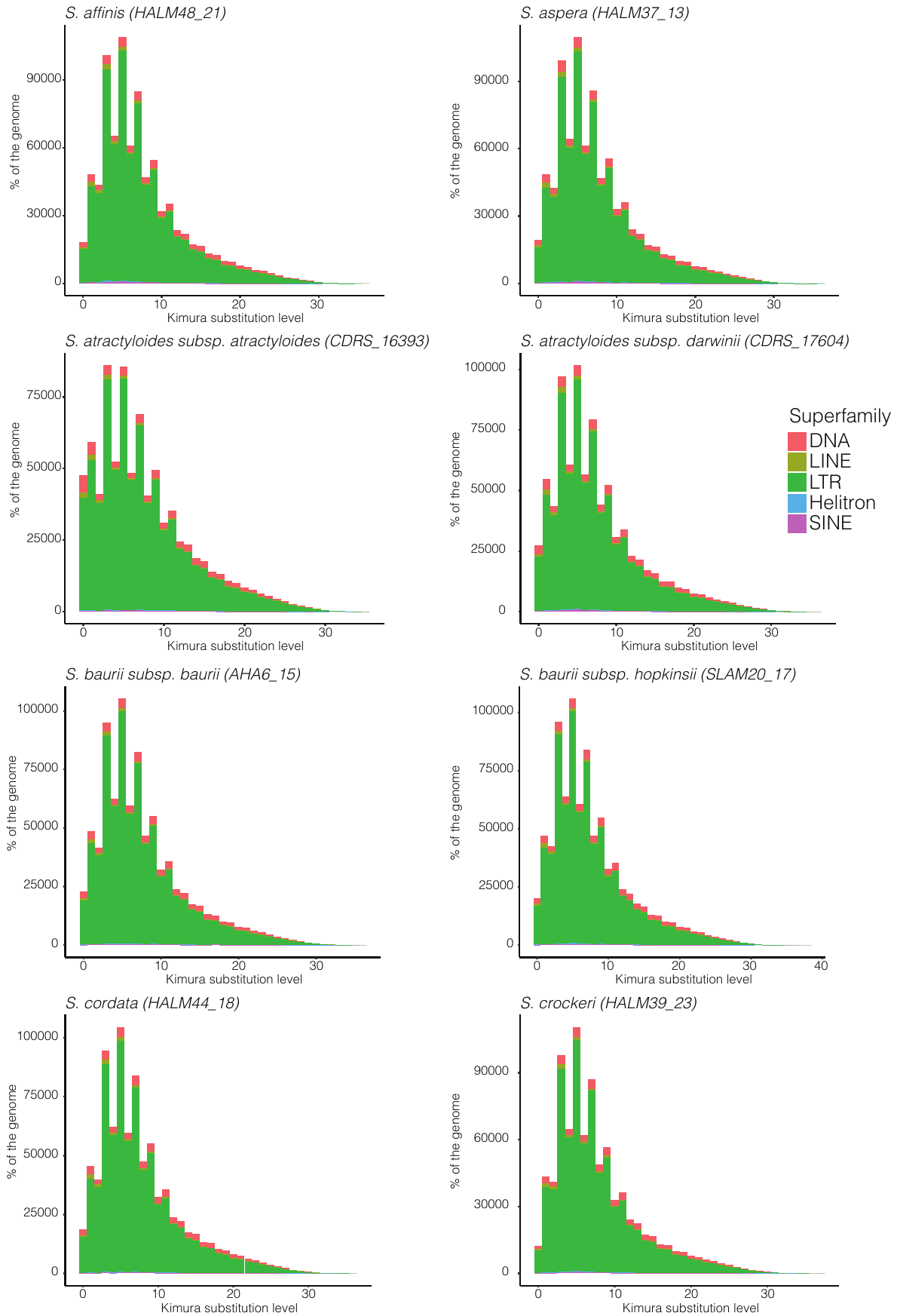

**Supplementary Figure 08** Repeat landscape plot showing the divergence of transposable elements (TEs) in the genome for *Scalesia* *divisa, S. gordilloi, S. helleri, S. incisa, S. microcephala, and S. pedunculata*. Different TE groups are represented by different colors.

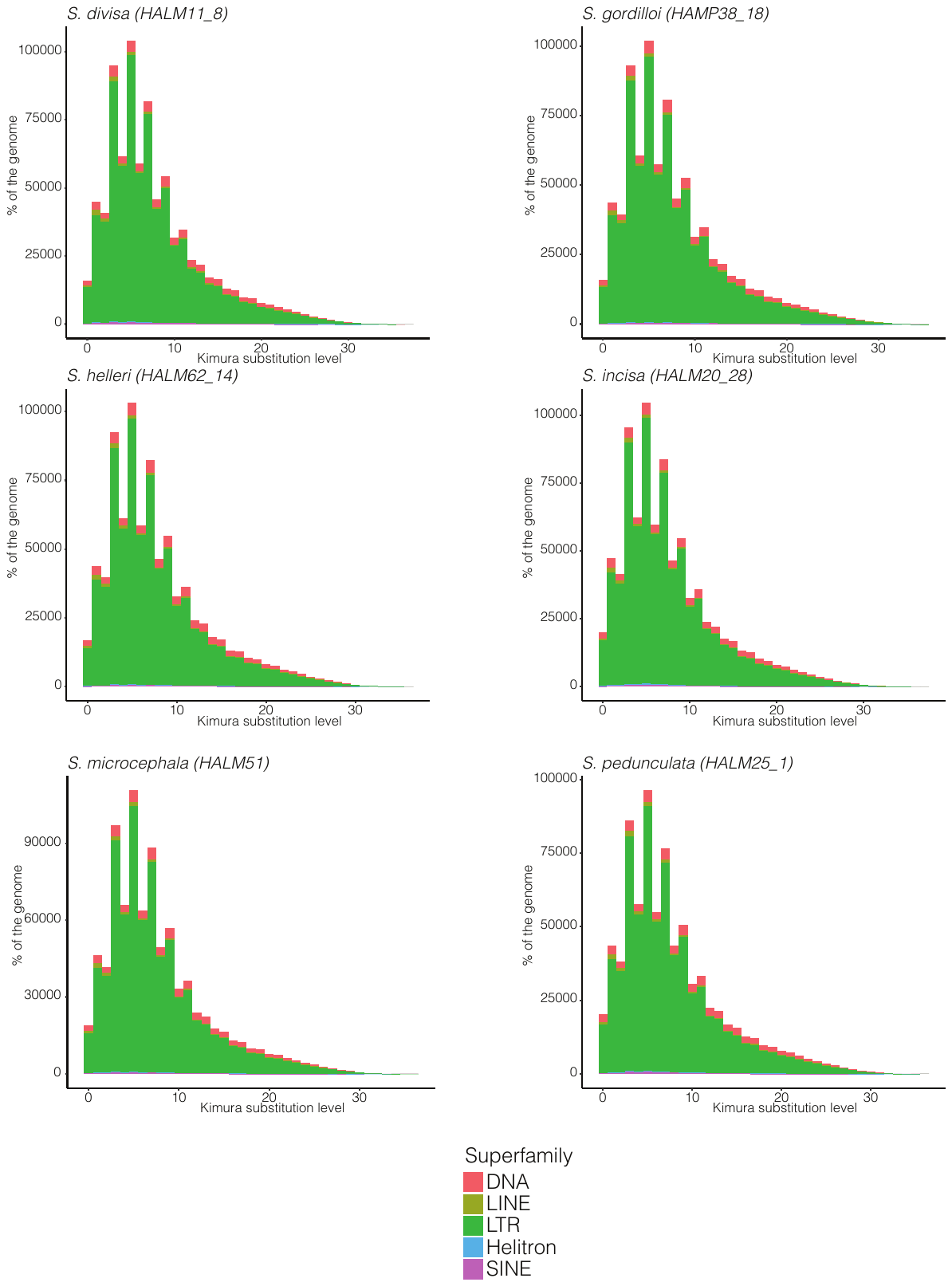

**Supplementary Figure 09** Repeat landscape plot showing the divergence of transposable elements (TEs) in the genome for *Scalesia* *retroflexa, S. stewartii, and S. villosa*. Different TE groups are represented by different colors.

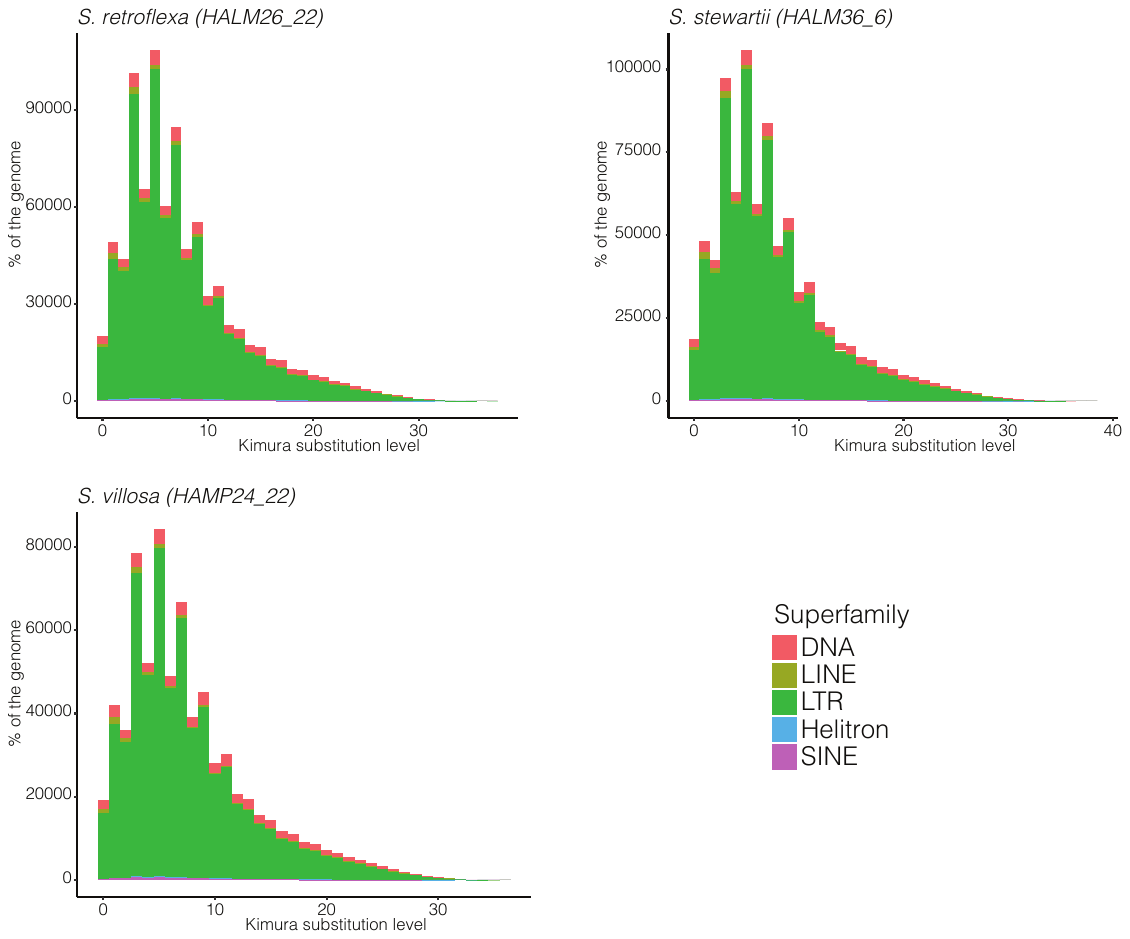

**Supplementary Figure 10** Repeat landscape plot showing the divergence of transposable elements (TEs) in the genome for *Scalesia* hybrids, including *S. incisa x divisa, S. crockeri x aspera, S. stewartii x atractyloides, S. crf retroflexa x helleri*. Different TE groups are represented by different colors.

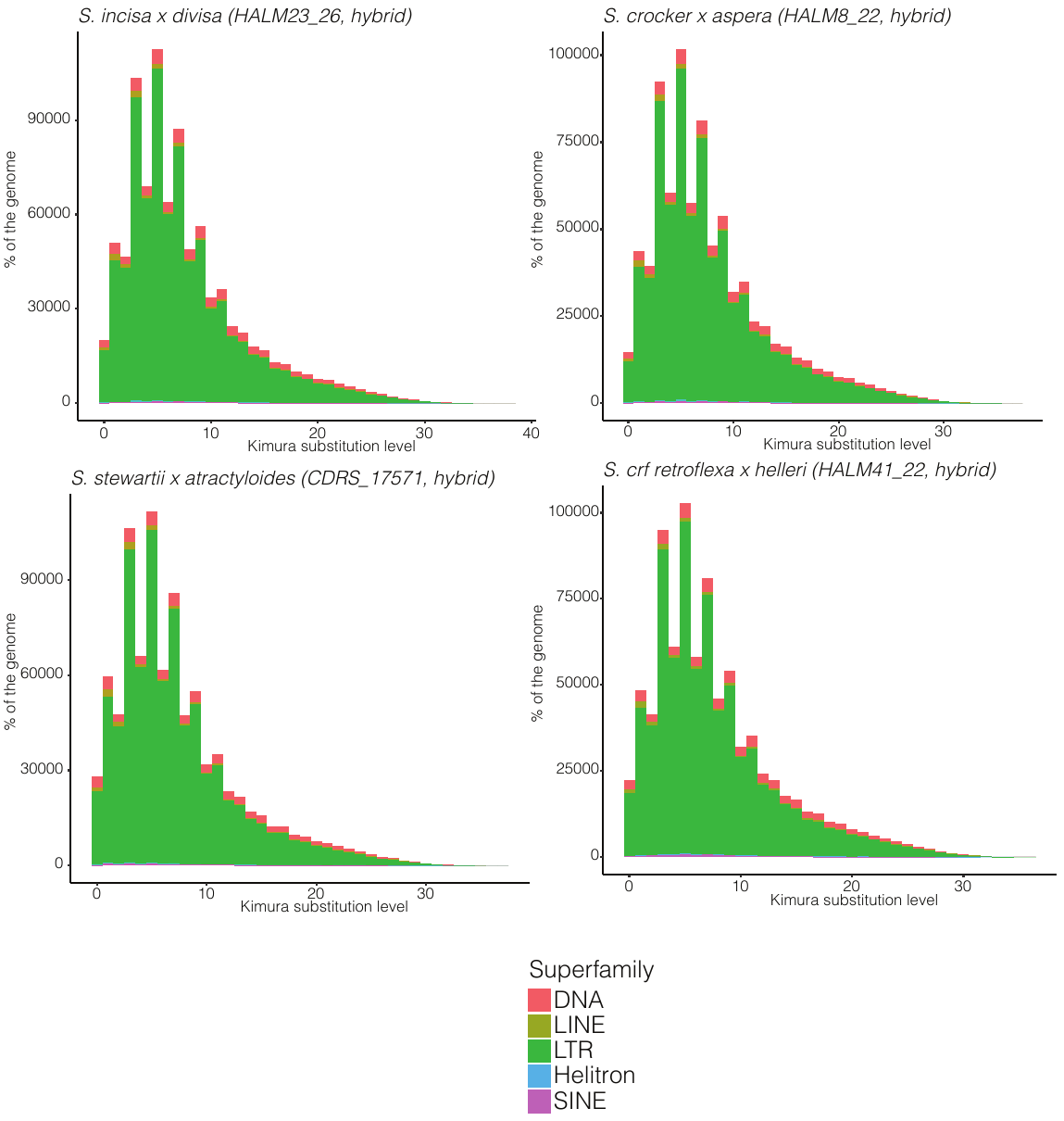

**Supplementary Figure 11** Upset plot of DNA elements, grouped in orthogroups, found within the Scalesia radiation. The number of orthogroups is plotted on the y-axis, and the x-axis shows the overlap between different genomes.

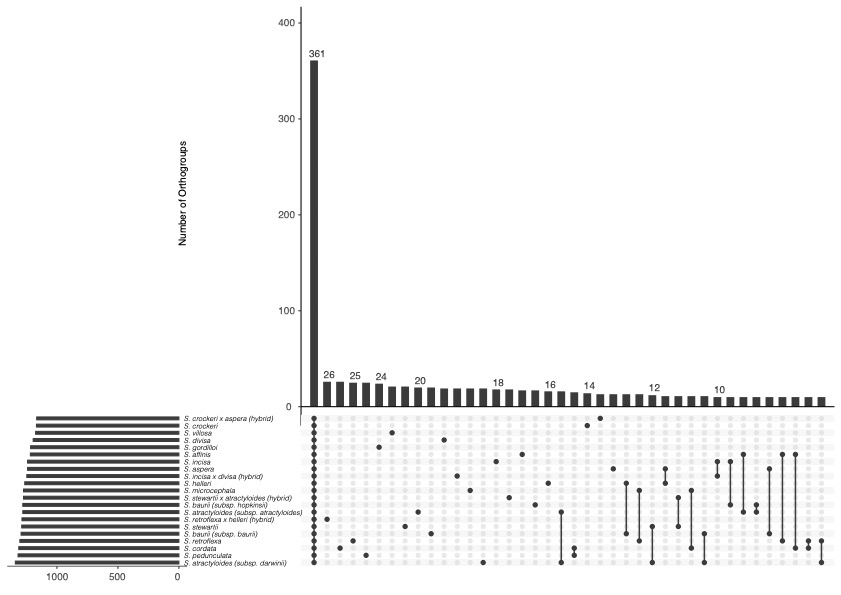

**Supplementary Figure 12** Upset plot of Helitrons, grouped in orthogroups, found within the Scalesia radiation. The number of orthogroups is plotted on the y-axis, and the x-axis shows the overlap between different genomes.

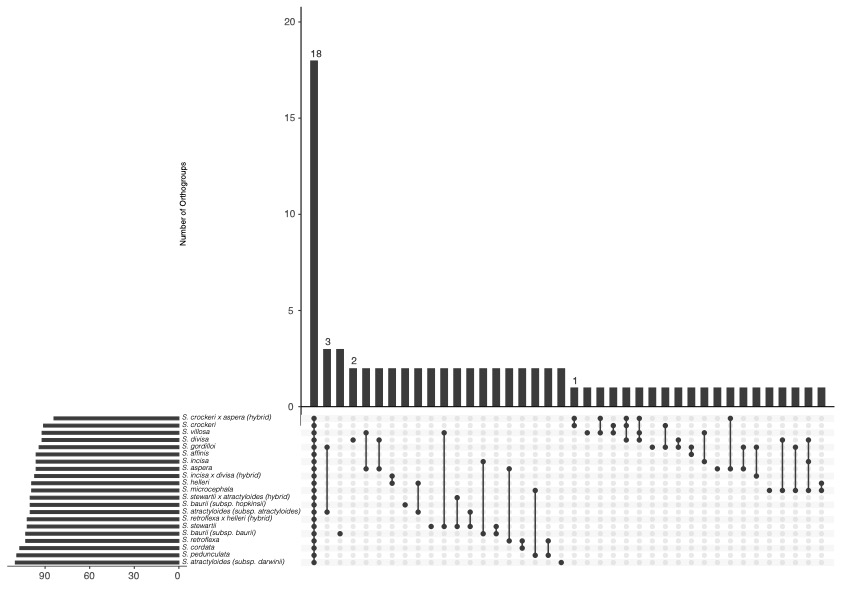

**Supplementary Figure 13** Upset plot of simple repeat elements, grouped in orthogroups, found within the Scalesia radiation. The number of orthogroups is plotted on the y-axis, and the x-axis shows the overlap between different genomes.

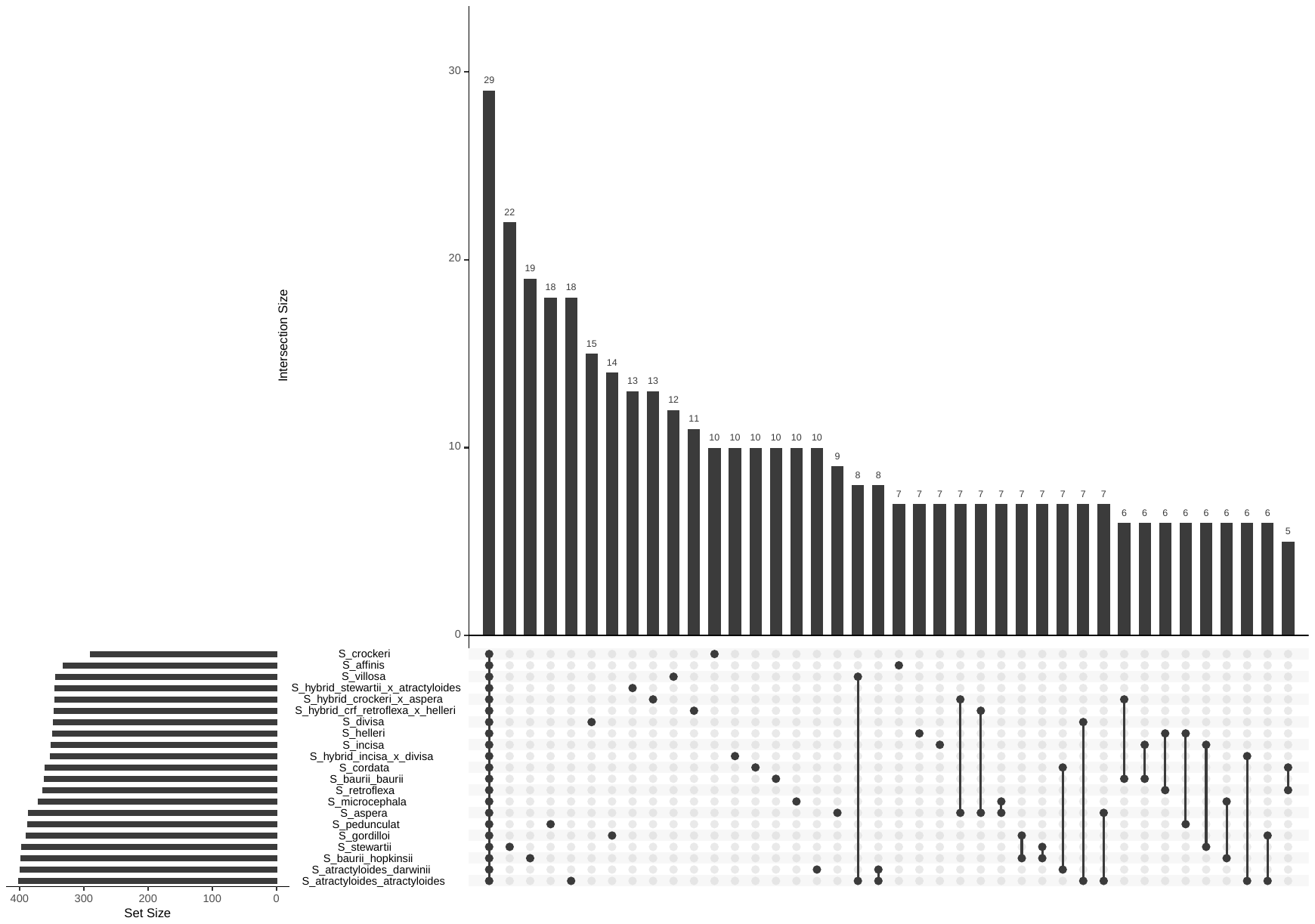

**Supplementary Figure 14** Upset plot of SINE, grouped in orthogroups, found within the Scalesia radiation. The number of orthogroups is plotted on the y-axis, and the x-axis shows the overlap between different genomes.

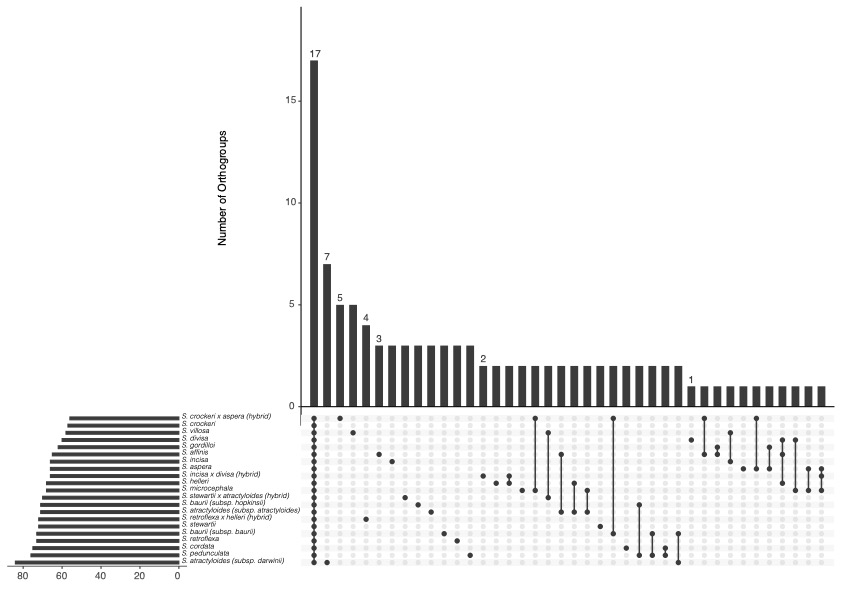

**Supplementary Figure 15** Upset plot of Satellite elements, grouped in orthogroups, found within the *Scalesia* radiation. The number of orthogroups is plotted on the y-axis, and the x-axis shows the overlap between different genomes.

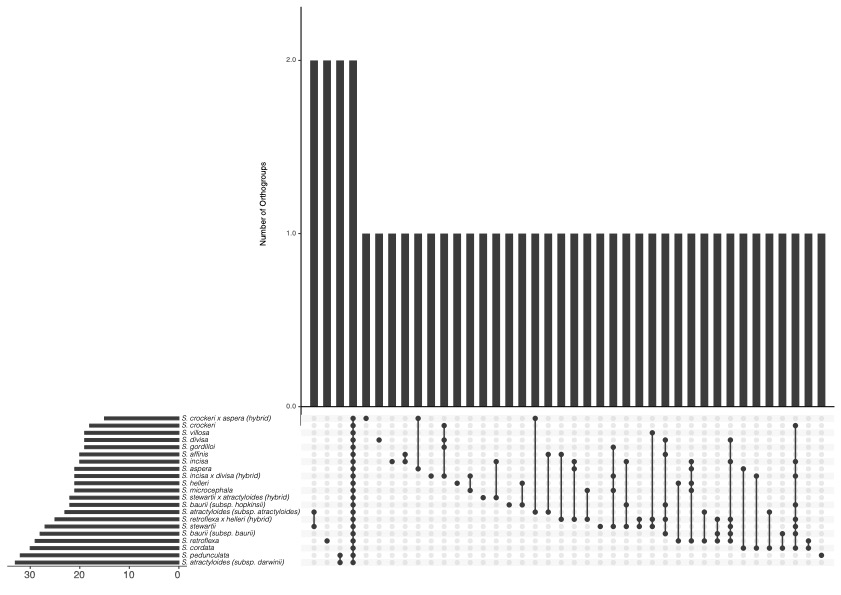

**Supplementary Figure 16** - TE Superfamily counts for the outgroup taxa. The count scale is separate for LTR/Copia and LTR/Gypsy due to the difference in scale.

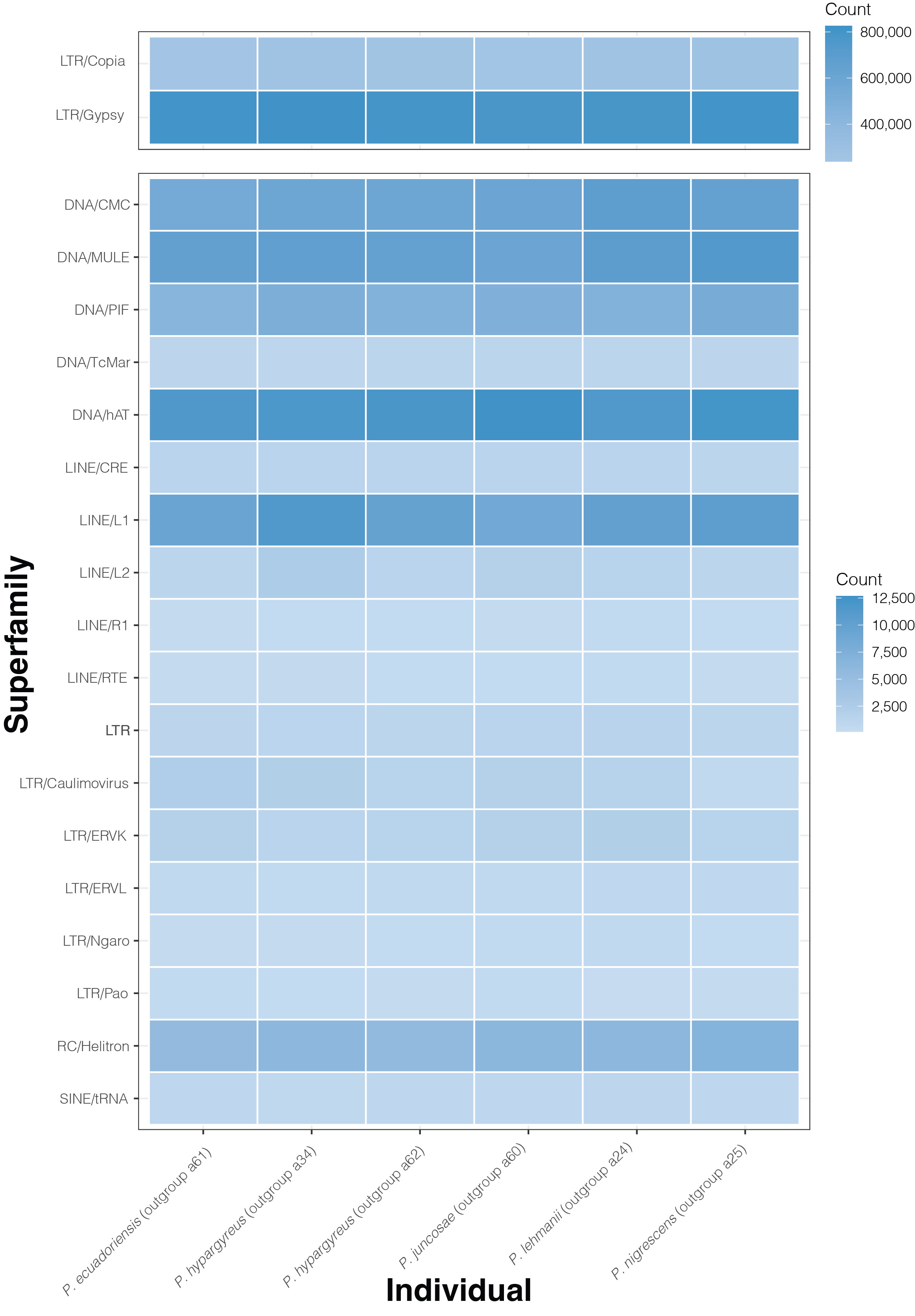

**Supplementary Figure 17** TEs and repeats 10kb upstream of genes (to the left) and within gene regions (to the right) for Helitron, Satellite, SINE, and Simple repeat elements. For each plot we show all *Scalesia* genes, leaf genes (important phenotypes), random parts of the genome, and candidate genes (with elevated dN/dS ratios, following Cerca et al 2022). Statistical significance were obtained with the Games−Howell test with a Holm–Bonferroni correction, as implemented in the R package ggstatsplot.

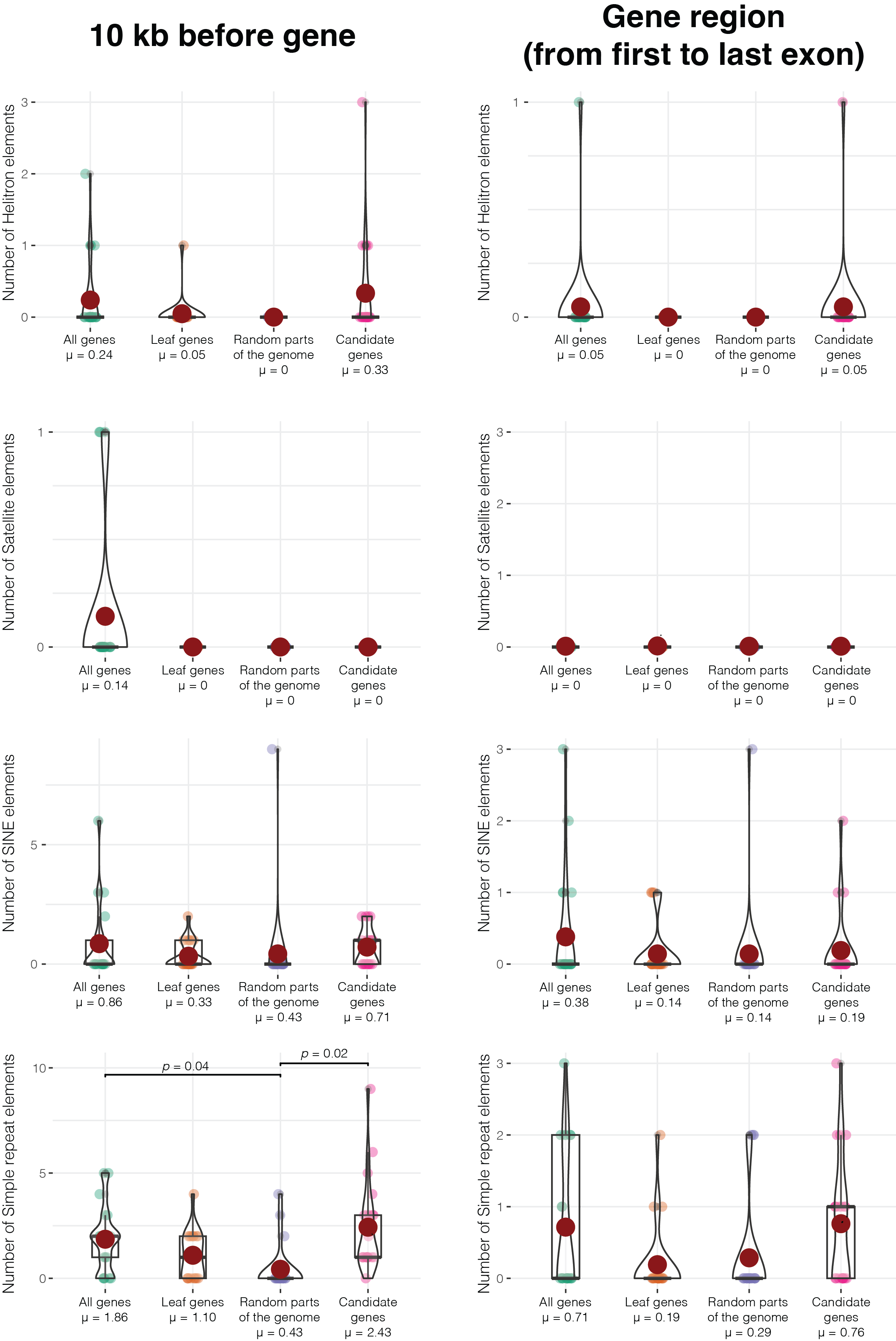
